# The RNA helicase DDX6 controls early mouse embryogenesis by repressing aberrant inhibition of BMP signaling through miRNA-mediated gene silencing

**DOI:** 10.1101/2021.11.29.470397

**Authors:** Jessica Kim, Masafumi Muraoka, Rieko Ajima, Hajime Okada, Atsushi Toyoda, Hiroshi Mori, Yumiko Saga

## Abstract

The evolutionarily conserved RNA helicase DDX6 is a central player of post-transcriptional regulation, but its role during embryogenesis remains elusive. We here demonstrated that DDX6 enables proper cell lineage specification from pluripotent cells by analyzing *Ddx6* KO mouse embryos and *in vitro* epiblast-like cell (EpiLC) induction system. Our study unveiled a great impact of DDX6-mediated RNA regulation on signaling pathways. Deletion of *Ddx6* caused the aberrant transcriptional upregulation of the negative regulators of BMP signaling, which accompanied with enhanced Nodal signaling. *Ddx6*^△^*^/^*^△^ pluripotent cells acquired higher pluripotency with a strong inclination toward neural lineage commitment. During gastrulation, abnormally expanded *Nodal* expression in the primitive streak likely promoted endoderm cell fate specification while inhibiting mesoderm development. We further clarified the mechanism how DDX6 regulates cell fate determination of pluripotent cells by genetically dissecting major DDX6 pathways: processing body (P-body) formation, translational repression, mRNA decay, and miRNA-mediated silencing. P-body-related functions were dispensable, but the miRNA pathway was essential for the DDX6 function. DDX6 may prevent aberrant transcriptional upregulation of the negative regulators of BMP signaling by repressing translation of certain transcription factors through the interaction with miRNA-induced silencing complexes (miRISCs). Overall, this delineates how DDX6 affects development of the three primary germ layers during early mouse embryogenesis and the underlying mechanism of DDX6 function.

**Author summary:** Gene expression occurs through the two steps: transcription (DNA to RNA) and translation (RNA to protein). Cells have very sophisticated regulatory processes working on various levels for the accurate gene expression. Post-transcriptional regulation, which includes all RNA-related controls, is crucial because it enables fine-tuning and rapid alteration of gene expression. RNA-binding proteins and non-coding RNAs are the two main players of post-transcriptional regulation. DDX6, the subject of our study, is an RNA-binding protein, more specifically an RNA helicase, which can unwind or rearrange RNA secondary structures. Its diverse molecular and cellular functions have been reported, but its embryogenic role is unknown. Here, we describe DDX6 function during early mouse embryogenesis and the underlying mechanism using genetic methodology. DDX6 enables proper cell lineage specification of pluripotent stem cells by mainly regulating BMP signaling through miRNA-mediated gene silencing. As DDX6- mediated RNA regulation affected signaling pathways, the loss of *Ddx6* had a wide impact on developmental processes from pluripotency to embryo patterning. In addition, we identified which DDX6 molecular function is essential during early embryogenesis by genetically dissecting its main pathways.

## Introduction

Post-transcriptional regulation, located in the middle layer of gene expression, is a critical controlling point where many mRNA regulatory processes occur. RNA-binding proteins (RBPs) and non-coding RNAs are the two main players (Kaikkonen et al., 2011; Dassi, 2017). Among diverse RBPs, RNA helicases are characterized by their wide range of involvement in RNA metabolism by binding RNA or remodeling ribonucleoprotein complexes (RNPs) (Tanner & Linder, 2001; Jankowsky, 2011). DEAD box proteins compose the largest RNA helicase family sharing the Asp-Glu-Ala-Asp (DEAD) motif and having ATP-dependent RNA unwinding activity (Linder & Jankowsky, 2011). Mice have 43 DEAD box RNA helicases. Among them, we focused on DDX6, an evolutionarily conserved throughout eukaryotes (Weston & Sommerville, 2006). DDX6 participates in many aspects of RNA metabolism: processing body (P-body) formation (Serman et al., 2007; Minshall et al., 2009), stress granule assembly, mRNA storage, mRNA decay (Coller et al., 2001), translational repression (Coller & Parker, 2005; Kamenska et al., 2016), microRNA (miRNA) pathway (Chu and Rana et al., 2006; Eulalio et al., 2007), and translational promotion (Scheller et al., 2009; Wang et al., 2015). Many molecular and cellular studies on DDX6 have been reported, but investigation of its embryonic developmental function is limited.

Gastrulation is a milestone in embryogenesis because the primary germ layers that give rise to all cell types develop during this developmental event. The three germ layers of the embryo, the ectoderm, mesoderm, and endoderm, basically originate from the inner cell mass (ICM) of the blastocyst. Pluripotent embryonic stem cells (ESCs) can be derived from the ICM of E3.5 early blastocysts or the epiblast of E4.5 late blastocysts (Evans & Kaufman, 1981; Thomson et al., 1995; Brook & Gardner, 1997; Thomson et al., 1998). Cells at these stages are in the naive (ground) pluripotent state. Soon after implantation, naive epiblasts differentiate into the primed pluripotent state in which cells become capable of committing to a certain lineage during gastrulation (Nichols & Smith, 2009; Smith, 2017; Mulas et al., 2017). The recently developed *in vitro* model, epiblast-like cells (EpiLCs) permit more precise staging of mouse pluripotent cells. Transcriptomic analysis demonstrated that EpiLCs, induced from ESCs, have similar properties to the epiblast of post-implanted, pre-gastrulating (E5.5∼6.0) embryos (Hayashi et al., 2011).

These *in vitro* systems enable detailed examination of pre- and early post-implantation embryos which are normally difficult to investigate *in vivo* due to their small size.

From the 8-cell stage, transcription factors and signaling pathways play a major role in cell fate determination (Rossant, 2018). The transforming growth factor-β (TGF-β) superfamily is one of major signaling pathways involved in early mammalian development. Bone morphogenetic protein (BMP) and Nodal belong to a different subgroup and they utilize distinctive serine/threonine kinase receptors for signal transduction. Activated receptors phosphorylate downstream intracellular mediators, receptor SMADs 1, 5 and 8 for BMP signaling, and SMADs 2 and 3 for Nodal signaling (Liu et al., 1996; Massague, 1996). BMP and Nodal signaling are known to mutually antagonize each other, and this antagonism mainly occurs intracellularly through the competition for the common signal mediator SMAD4 (Candia et al., 1997; Furtado et al., 2008; Katsu et al., 2013; Soh et al., 2020). BMP signaling has multiple roles during early post-implantation development. It is required for extra-embryonic mesoderm formation (Zhang & Bradley, 1996), primordial germ cell (PGC) induction (Lawson et al., 1999), mesoderm development and patterning (Winnier et al., 1995), and inhibiting premature neural differentiation (Di-Gregorio et al., 2007). BMP signaling is dispensable for ESC self-renewal, but required for proper differentiation (Fei et al., 2010; Morikawa et al., 2016). Nodal pathway exerts its influence when ESCs undergo transition from the naive pluripotency. Nodal is important for securing primed pluripotency with the capacity to differentiate into multi-lineages (Mulas et al., 2017).

There are two previous studies that assessed the role of DDX6 in mouse and human pluripotent cells. In mESCs, DDX6 is necessary for translational repression of miRNA targets and *Ddx6* knockout (KO) cells exhibited similar phenotypes to *Dgcr8* KO ESCs, which lack all miRNAs (Freimer et al., 2018). Another study elucidated the relationship between stem cell potency and P-body-mediated translational regulation. As an essential factor of P-body formation, once DDX6 was depleted, P-bodies were disassembled then translationally suppressed target mRNAs, including many transcription factors and chromatin regulators, re-entered the translation pool. The resultant increased expression of target genes altered chromatin organization, and made both human and mouse primed ESCs become more naively pluripotent by being resistant to differentiation (Di Stefano et al., 2019).

However, these examinations were conducted on ESCs or very early differentiation stages. To deepen our understanding of the role of DDX6 as a key post-transcriptional regulator during early mouse embryogenesis, we examined *Ddx6* KO embryos. Gastrulation stages were investigated in embryos and the earlier time points were assessed using an ESC-to-EpiLC induction model. This study revealed that DDX6 exerts potent effects on the development of the three primary germ layers by preventing aberrant inhibition of the BMP signaling pathway.

Furthermore, through genetic dissection of the DDX6 pathways, we found that DDX6 works through the miRNA pathway, while P-bodies are dispensable during early development.

## Results

### *Ddx6* knockout results in embryonic lethality with severe morphological defects

Before investigating the functions of DDX6 during embryogenesis, we examined its expression pattern through DDX6 immunohistochemistry (IHC). In embryonic day (E) 6.5 embryos, DDX6 was highly and ubiquitously expressed as forming cytoplasmic foci (Fig. S1A). All DDX6 foci co-localized with DCP1A foci, a P-body specific marker, indicating that DDX6 is expressed in P-bodies. At E7.5, DDX6 expression was strongest in the epiblast and there were many clear P-body foci (Fig. S1B). BRACHYURY-positive emerging and migrating mesoderm cells had relatively weaker DDX6 expression and a fewer number of P-bodies. In E8.5 embryos, DDX6 expression was observed in all areas including the neuroepithelium, the tailbud, and somites (Fig. S1C). DDX6 is ubiquitously expressed in early embryos in P-bodies.

To clarify the role of DDX6 in embryonic development, we generated a *Ddx6*^△^*^/+^* mouse line and crossed heterozygous mice to get *Ddx6*^△^*^/^*^△^ (KO) embryos. All *Ddx6*^△^*^/^*^△^ embryos died by E11.5 (Table 1), and developmental defects were already observed from E6.5 (Fig. 1A). At E7.5, mutants were smaller, but an extraembryonic body part was developed and embryos formed a cylindrical shape. Morphological defects became prominent from E8.5, and more phenotypical variances appeared (Fig. S2A). E8.5 *Ddx6* KO embryos were categorized into three groups according to the severity of posterior body defects (Fig. 1A). Type I mutants developed clear head folds with a short posterior body part. Type II embryos had a head fold structure but mid-posterior body development was more disrupted than Type I. Type III were tiny and were unable to fully escape the egg cylinder shape. The frequency of each mutant was estimated as Type III (∼60%), Type II (∼30%), and Type I (∼10%). From E9.5, *Ddx6*^△^*^/^*^△^ embryos were divided into two groups: one that developed some mid-posterior body and another with marked posterior truncation (Fig. S2B). Therefore, DDX6 is necessary for mouse embryonic development.

**Table 1.** Antibodies.

| Antigen | Usage | Concentration | Manufacturer | Reference # |
| --- | --- | --- | --- | --- |
| **Rck/p54**<br>**(DDX6)** | IHC, ICC | 1:300 | MBL | PD009 |
| **DCP1A** | IHC, ICC | 1:200 | ABNOVA | H00055802-M06 |
| **BRACHYURY** | IHC | 1:400 | R&D SYSTEMS | O15178 |
| **NANOG** | IHC | 1:200 | NOVUS<br>BIOLOGICALS | NB100-58842 |
| **TuJ1** | IHC | 1:1500 | Abcam | Ab18207 |
| **SOX1** | IHC | 1:100 | NOVUS<br>BIOLOGICALS | AF3369 |
| **SOX2 (Y-17)** | IHC | 1:200 | SANTA CRUZ<br>BIOTECHNOLOGY | sc-17320 |
| **DCX** | IHC | 1:400 | Cell Signaling<br>TECHNOLOGY | (A8L1U) 14802 |

**Figure 1.**
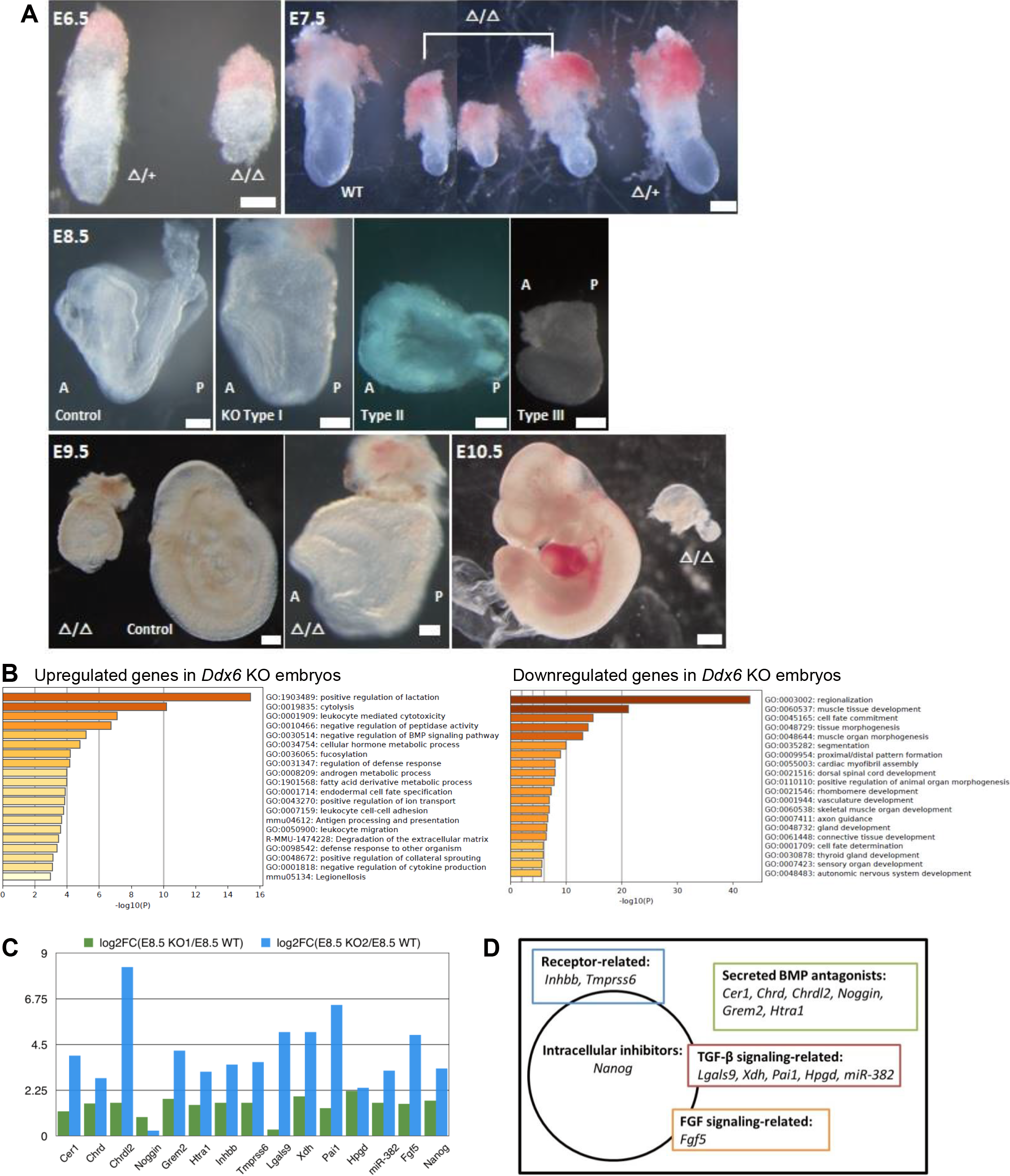
Characterization of *Ddx6*^△^*^/^*^△^ embryos. (A) *Ddx6*^△^*^/^*^△^ embryos exhibit growth delay and morphological defects. E6.5 (scale: 100 μm), E7.5 & E8.5 (200 μm), E9.5 (100 μm for group; 200 μm for KO), and E10.5 (500 μm). (B-D) Gene expression analyses by RNA-seq. (B) Gene ontology (GO) term enrichment analysis of most upregulated and downregulated genes. (C) RNA-seq data comparing expression of negative regulators of the BMP pathway in E8.5 *Ddx6* KOs with that in E8.5 WT. (D) Classification of types of the upregulated negative regulators of BMP signaling.

### Transcriptomic analyses revealed negative regulation of BMP signaling in *Ddx6*^△^*^/^*^△^ embryos

To find possible causes of defects, we performed RNA sequencing (RNA-seq). Two E8.5 *Ddx6* KO cDNA libraries were generated. Being consistent with morphological phenotypes, gene ontology (GO) term enrichment analysis indicated that genes of major developmental processes, especially the formation of mesoderm derivatives, were downregulated in *Ddx6* KO libraries (Fig. 1B). In contrast, the terms associated with cell death, immune response, cell metabolism, and negative regulation of BMP signaling pathway-related genes were highly upregulated. Negative regulation of the BMP signaling pathway was notable, because BMP signaling has multiple important roles during early embryogenesis. Various kinds of BMP negative regulators were upregulated in *Ddx6*^△^*^/^*^△^ embryos (Fig. 1C). Based on the reported functions, genes that are listed as the negative regulators of BMP signaling were classified into five clusters: receptor-related (*Inhbb,* and *Tmprss6*), secreted BMP antagonists (*Cer1, Chrd, Chrdl2, Noggin, Grem2,* and *Htra1*), TGF-β signaling-related (*Lgals9, Xdh, Pai1, Hpgd,* and *miR-382*), FGF signaling-related (Fgf5), and intracellular inhibitor (*Nanog*) (Fig. 1D). As the genes that are related to the inhibition of BMP signaling were highly upregulated, we examined whether BMP signaling is repressed in *Ddx6*^△^*^/^*^△^ embryos. We reasoned that if BMP signal transduction is indeed dysfunctional, then *Ddx6*^△^*^/^*^△^ embryos would exhibit the representative phenotypes of BMP signaling mutant embryos.

### *Ddx6*^△^*^/^*^△^ embryos display phenotypes arising from the disrupted BMP signaling pathway

#### 1) Mesoderm formation defects

BMP signaling is required for mesoderm formation and posterior body development (Winnier et al., 1995; Reversade et al., 2005). Whole-mount *in situ* hybridization (WISH) with *Otx2* probe, which marks a head region, indicated the lack of posterior body in Type III mutants (Fig. 2A). We analyzed mesodermal defects using E8.5 embryo RNA-seq data. Two KO samples showed difference in mesoderm-related gene expression (Fig. 2B-C). This difference was consistent with the types of mutants that compose each library. KO1, constituted with one Type II and one Type III mutant, had similar expression pattern to WT. In contrast, KO2, constituted with three Type III mutants, had a very different transcriptome, in which the expression of early mesoderm marker genes was higher, but that of differentiating mesoderm was significantly downregulated. Visualization of *Brachyury (T)* expression via WISH revealed that the primitive streak formed in all types of E8.5 *Ddx6*^△^*^/^*^△^, but somites were barely developed (Fig. 2D).

**Figure 2.**
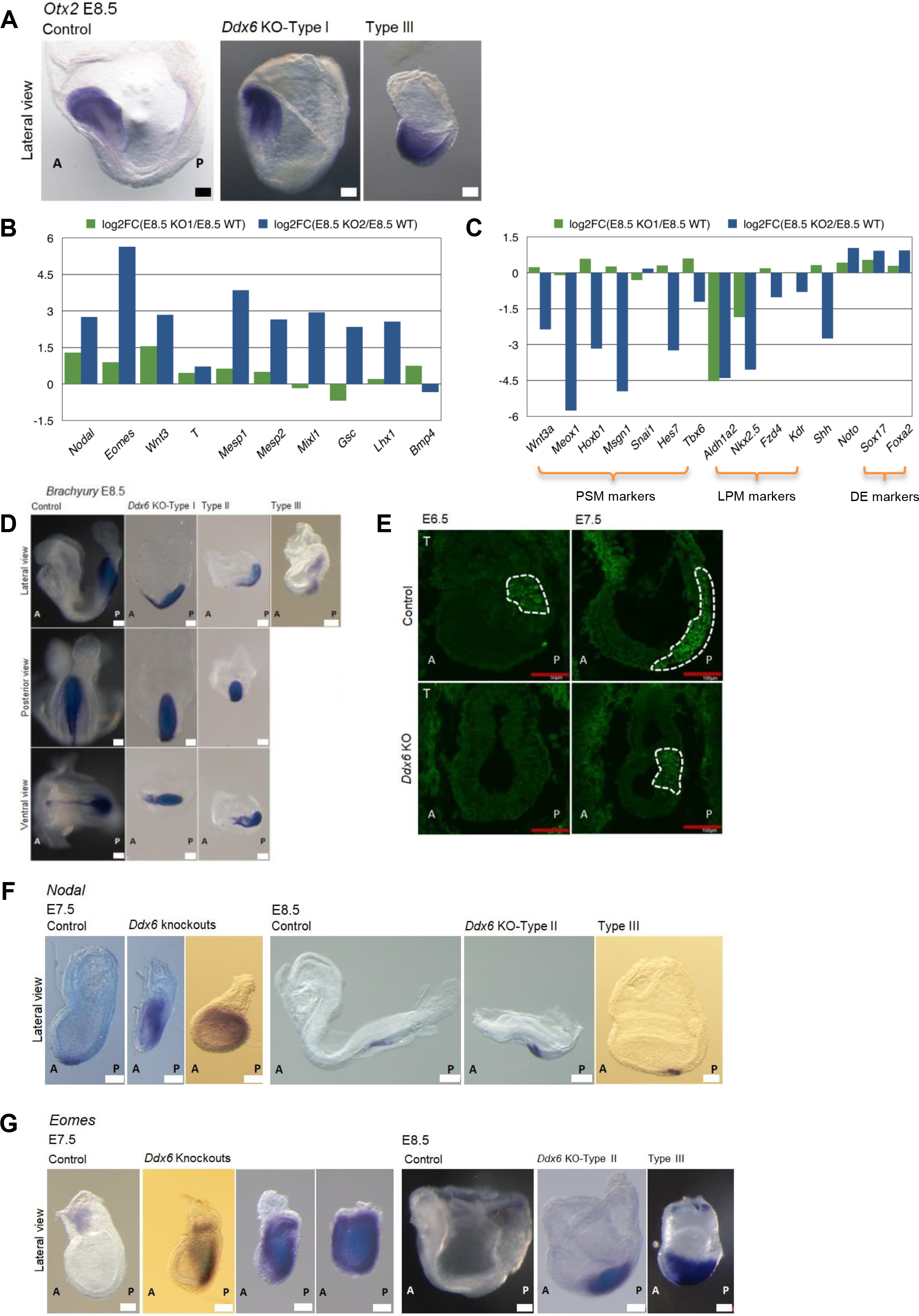
*Ddx6***^△^***^/^***^△^** embryos have defects in mesoderm development. (A) Whole-mount ISH of E8.5 embryos with an *Otx2* probe (100μm, n = 3). (B-C) RNA-seq data comparing expression of several key genes in E8.5 *Ddx6* KOs with that in E8.5 WT. (B) primitive streak and early mesoderm-related genes. (C) differentiated mesoderm and endoderm markers. PSM: paraxial mesoderm, LPM: lateral plate mesoderm, DE: definitive endoderm. (D) Whole-mount ISH of E8.5 embryos with a *Brachyury* probe (100 μm, n = 5). (E) E6.5 & E7.5 embryo frozen section IHC for T (50 μm for E6.5, n = 2; 100 μm for E7.5, n = 3). (F) Whole-mount ISH of E7.5 & E8.5 embryos with a *Nodal* probe (scale: 100 μm for E7.5 & E8.5 Type III; 200 μm for E8.5 Control & Type II, n = 4 for E7.5, n = 3 for E8.5). (G) Whole-mount ISH of E7.5 & E8.5 embryos with an *Eomes* probe (100 μm, n = 3 for each time point).

Gastrulation begins ∼E6.5 as the primitive streak forms and is preceded by *Brachyury* expression (Snell and Stevens, 1966; Rivera-Perez and Magnuson, 2005). Unlike WT, *Ddx6*^△^*^/^*^△^ embryos started expressing BRACHYURY (T) from E7.5 (Fig. 2E). Therefore, there is a delay of primitive streak formation in *Ddx6*^△^*^/^*^△^ embryos, and their shortened and widened primitive streak suggests that the nascent mesoderm population has defects in differentiation and subsequent ingression.

An evidence of suppressed BMP signaling in *Ddx6*^△^*^/^*^△^ embryos was increased Nodal signaling. BMP and Nodal signaling are often in a competitive relationship, and they can suppress each other. During gastrulation, a gradient of NODAL activity patterns the primitive streak and allocates mesendoderm progenitors. NODAL and its downstream target EOMES together define the anterior primitive streak (APS), from which cardiac mesoderm and definitive endoderm progenitors are specified (Brennan et al., 2001; Arnold et al., 2008; Teo et al., 2011; Costello et al., 2011). WISH showed that *Nodal* expression was confined to the node in E7.5 WT embryos, but its expression was highly spread over the proximal-posterior region in *Ddx6*^△^*^/^*^△^ embryos (Fig. 2F). The expression was eventually restricted to the node by E8.5, but the expression level remained high. *Eomes* is highly expressed in the extraembryonic ectoderm and the posterior part of the epiblast at E6.5. Its expression moves distally to the primitive streak at E7.5 (Russ et al., 2000), and is reduced by E8.5 in the WT embryos. However, E8.5 *Ddx6*^△^*^/^*^△^ embryos retained high-level expression (Fig. 2G). There were two types of knockout embryos at E7.5, which could reflect the difference in severity of mutant phenotypes. One type had a slightly stronger expression level of *Eomes,* but the positive area was similar to E6.5 WT. The other had strong expression encompassing nearly the entire body (Fig. 2G). RNA-seq analyses showed that the KO2 sample had downregulated expression of differentiated mesoderm genes, while the expression of endoderm lineage genes was upregulated. KO2 exhibited higher expression of mesendoderm progenitor markers (*Mixl1,* and *Gsc*), endoderm progenitor marker (*Lhx1*), and definitive endoderm markers (*Sox17,* and *Foxa2*) (Fig. 2B-C). The ‘Endoderm cell fate specification’ category was also enriched in GO term analysis of most upregulated genes in *Ddx6*^△^*^/^*^△^ (Fig. 1B). Therefore, posteriorly expanded high expression of the APS marker *Nodal* and *Eomes* disturbed the patterning of the primitive streak, which likely directed mesendoderm progenitors toward the endodermal lineage. Transcript levels of some key genes were further assessed by RT-qPCR using separately prepared Type III mutant embryos and the results were consistent with the RNA-seq data (Fig. S2C). Altogether, *Ddx6*^△^*^/^*^△^ embryos have defects in posterior body development and mesoderm differentiation like as BMP signaling mutant embryos.

#### 2) Premature neural induction

BMP signaling also prevents the premature neural induction (Di-Gregorio et al., 2007). We examined whether this function was also impaired in *Ddx6*^△^*^/^*^△^ embryos. SOX1 is the earliest neuroectoderm marker, and it is normally not detected until E7.5 in WT embryos (Wood & Episkopou, 1999; Di-Gregorio et al., 2007). However, E6.5 *Ddx6*^△^*^/^*^△^ embryos exhibited clear SOX1 expression in all epiblast cells (Fig. 3A). Additionally, RNA-seq found premature neuronal differentiation in *Ddx6*^△^*^/^*^△^. The markers of neural stem cells (NSCs) and neural progenitor cells (NPCs), such as *Sox1* and *Pax6,* were downregulated, but genes of neuron-restricted progenitors and differentiated post-mitotic neuronal cells were upregulated (Fig. 3B- C). Section IHC confirmed that protein expression was similar to transcript levels. The earliest neuroectoderm marker, SOX1, and a persistent marker of NSC and NPC, SOX2 (Ellis et al., 2004), were weakly expressed in E8.5 *Ddx6*^△^*^/^*^△^ embryos (Fig. 4D). DCX is a marker of neuronal precursors or early immature neurons, and is expressed in migrating neurons. The uppermost part of the cortical plate, which is composed of the most recently migrated neurons, also exhibits strong DCX immunoreactivity (Gleeson et al., 1999). In KO embryos, the intensity of DCX signal was stronger throughout the body. A strong DCX-positive layer was also observed (marked by a yellow arrow) (Fig. 3E). In summary, in *Ddx6*^△^*^/^*^△^ embryos, the neural lineage was precociously induced like in BMP receptor mutant embryos (Di-Gregorio et al., 2007). Moreover, *Ddx6*-deficient NSCs showed defects in maintaining self-renewal and prematurely differentiated.

**Figure 3.**
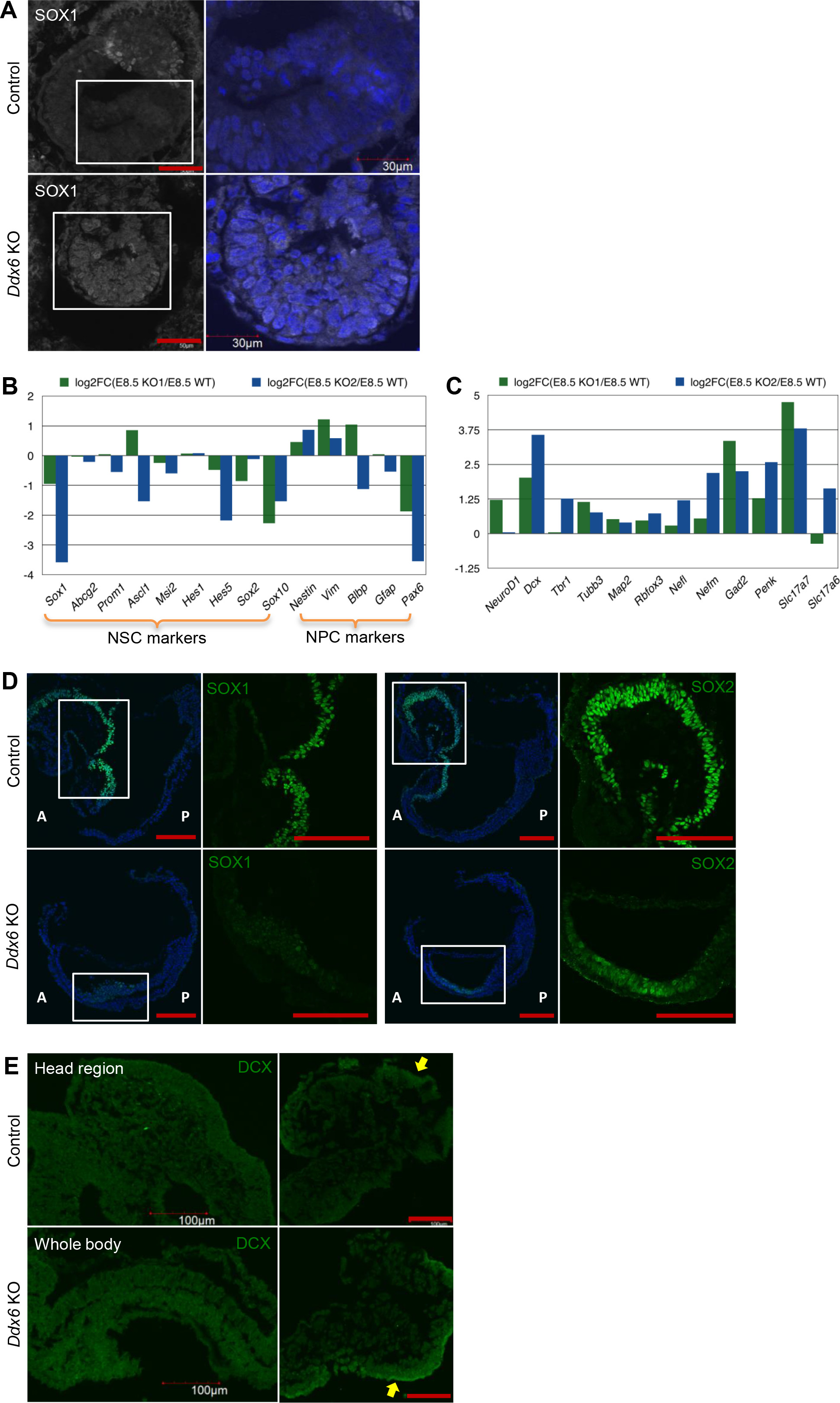
Precocious neural induction and premature differentiation are observed in *Ddx6*^△^*^/^*^△^ embryos. (A) E6.5 embryo frozen section IHC for SOX1 (50 μm for lower magnification, n = 2). (B-C) RNA-seq data comparing expression of several key genes in E8.5 *Ddx6* KOs with that in E8.5 WT. (B) NSC and radial glial cell (NPC) markers. (C) genes related to neuron-restricted intermediate progenitor & differentiated neuron. (D) E8.5 embryo frozen section IHC for SOX1 & SOX2 (scale: 100 μm, n = 3). (E) E8.5 embryo frozen section IHC for DCX (100 μm, n = 3).

**Figure 4.**
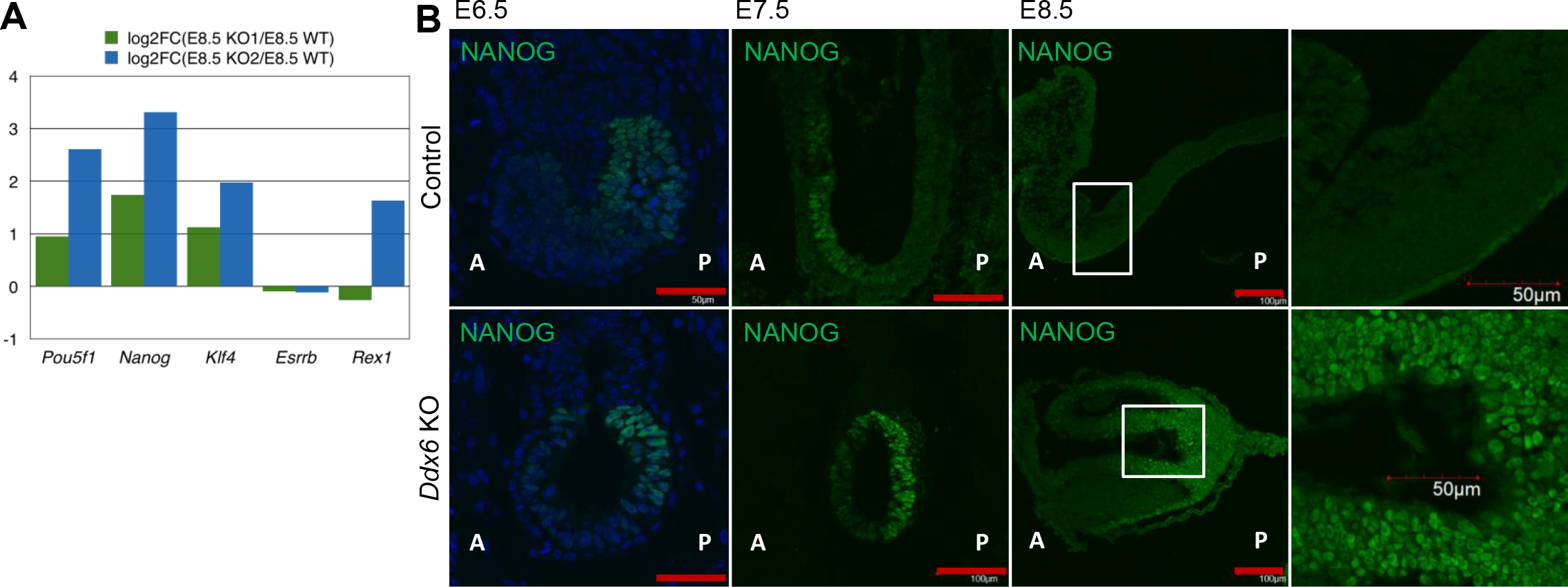
E8.5 *Ddx6***^△^***^/^***^△^** embryos retain strong naive and primed pluripotency. (A) RNA-seq data comparing expression of pluripotency marker genes in E8.5 *Ddx6* KOs with that in E8.5 WT. (B) E6.5∼E8.5 embryo frozen section IHC for NANOG (scale: 50 μm for E6.5, n = 2; 100 μm for E7.5, n = 3 for E7.5 & E8.5).

### Posterior epiblast of *Ddx6*^△^*^/^*^△^ embryos cannot exit the pluripotency on time

As described earlier, Nodal signaling, in an antagonistic relationship with BMP signaling, gets upregulated when BMP signaling is suppressed. One example was the increased expressions of *Nodal* and its downstream target *Eomes* in the primitive streak of E8.5 *Ddx6*^△^*^/^*^△^ embryos (Fig. 2F-G). Nodal signaling is also important for regulating primed pluripotency. Activin/Nodal signaling is required to induce *Nanog* transcription in mEpiSCs (Vallier et al., 2009). Therefore, we examined whether the expression of this another Nodal signaling target gene was also increased in *Ddx6*^△^*^/^*^△^ embryos. The core pluripotency factors *Nanog* and *Pou5f1* (*Oct4*) and a naive pluripotency marker, *Klf4,* were upregulated in *Ddx6*^△^*^/^*^△^ (Fig. 4A). However, the expression of another naive ground state marker, *Esrrb,* was similar to WT, and *Rex1* expression was higher only in the KO2. The partially retained expression of the naive pluripotency-specific genes in E8.5 embryos suggested that exit from the ground pluripotent state did not occur properly in *Ddx6* mutants. We examined the NANOG expression by section IHC (Fig. 4B). In WT embryos, NANOG expression was strongest in the primitive streak at E6.5, but at E7.5, its expression moved anteriorly and the posterior part was negative. Contrarily, E7.5 *Ddx6*^△^*^/^*^△^ embryos exhibited strong NANOG expression in the posterior epiblast. Ectopic and high expression of NANOG in the posterior part was maintained until E8.5 when its expression was not detected in the whole body of E8.5 WT embryos. Enriched GO terms among downregulated genes from RNA-seq, ‘Cell fate commitment’ and ‘Cell fate determination’ supported the strengthened primed pluripotency and the retained naive pluripotency of *Ddx6*^△^*^/^*^△^ embryos.

### *Ddx6*^△^*^/^*^△^ pluripotent cells also show repressed BMP signaling with enhanced Nodal signaling

We have examined E8.5 *Ddx6*^△^*^/^*^△^ embryos and considered repressed BMP signaling as a major cause of their developmental defects. We then looked for the earliest time point when inhibition of BMP signaling occurs in *Ddx6*^△^*^/^*^△^. There were no morphological abnormalities until E3.5 blastocysts and ESCs were successfully established from them. DDX6 was highly expressed in ESCs and EpiLCs in P-bodies, which were disassembled in *Ddx6*^△^*^/^*^△^ cells (Fig. S3A-B). The proliferation rate of *Ddx6*^△^*^/^*^△^ ESCs was lower (Fig. 5A), but they had no defects in maintaining pluripotency over many passages. Rather, like E8.5 *Ddx6*^△^*^/^*^△^ embryos, they expressed higher levels of pluripotency genes, such as *Oct4, Nanog, Sox2*, *Klf4,* and *Rex1,* than WT ESCs (Fig. 5B).

**Figure 5.**
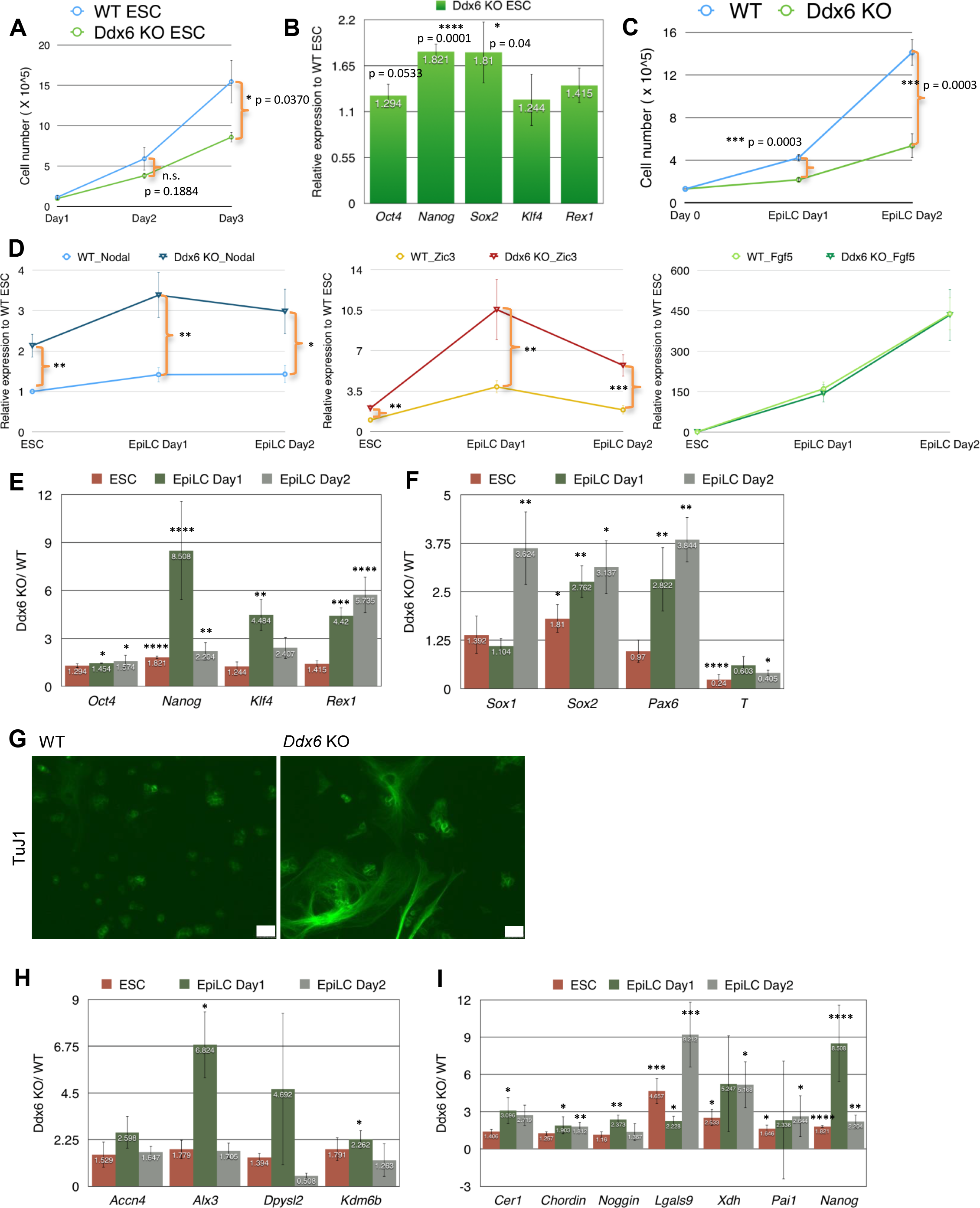
Characterization of *Ddx6***^△^***^/^***^△^** pluripotent cells. (A) Cell counting of ESCs over three-day culture period. Mean ± SEM. The statistical significance was calculated by Student’s t test (n = 5). (B) RT-qPCR examining relative expression of pluripotency markers in *Ddx6*^△^*^/^*^△^ ESCs to WT ESCs. Mean ± SEM. Student’s t test (n = 7∼9). (C) Cell counting during ESC-to-EpiLC induction period. Mean ± SEM. Student’s t test (n = 13 for WT, n = 7 for *Ddx6*^△^*^/^*^△^). (D) RT-qPCR examining the expression pattern of *Nodal*, *Fgf5* and *Zic3*. Mean ± SEM. Student’s t test (n = 9 for *Nodal* & *Fgf5*, n = 7 for *Zic3*) (*p ≤ 0.05, **p ≤ 0.01, ***p ≤ 0.001, ****p ≤ 0.0001). (E-F, H-I) RT-qPCR analysis of the expression trend of several key genes during EpiLC induction period. Each bar represents the relative expression of *Ddx6*^△^*^/^*^△^ cells to WT cells at the indicated time point. Mean ± SEM. Student’s t test. (E) major pluripotency genes (n = 7∼9). (F) early neuroectoderm and mesendoderm lineage markers (n = 7∼9). (H) BMP-SMAD1/5 target genes (n = 5). (I) the negative regulators of BMP signaling (n = 6∼9). (G) TuJ1 ICC on Day1 of monolayer differentiation (scale: 50 μm) (n = 3).

Since *Ddx6*^△^*^/^*^△^ blastocysts and ESCs did not exhibit notable abnormalities, we examined their next developmental capacity. We conducted ESC-to-EpiLC induction to mimic the natural *in vivo* development. The transition of the naive ground state ESCs to primed pluripotent state EpiLCs takes two days and EpiLC Day1 is regarded as a transition state that exhibits a distinctive open chromatin landscape and transcriptome (Yang et al., 2019). During EpiLC induction, the difference in cell number between WT and *Ddx6*^△^*^/^*^△^ cells markedly increased (Fig. 5C). We then investigated the expression pattern of several key genes during EpiLC induction by RT-qPCR. Firstly, we checked well-known EpiLC markers *Fgf5* and *Nodal,* and an important regulator of the transition state, ZIC3, which exhibits peak expression on EpiLC Day1 (Yang et al., 2019). *Ddx6*^△^*^/^*^△^ cells had significantly higher expression of *Nodal* and *Zic3*, but there was no difference in *Fgf5* expression (Fig. 5D). After confirming successful induction, we examined the expression profile of pluripotency and early differentiation-related genes. As shown in Fig. 5B, *Ddx6*^△^*^/^*^△^ ESCs exhibited slightly higher expression of pluripotency genes and this difference increased during EpiLC induction (Fig. 5E). They had much higher expression of naive pluripotency markers (*Klf4, Rex1*), suggesting that even though an overall transition was made to the EpiLC state, cells failed to completely exit from the ground state. We also noted a change in differentiation capacity of *Ddx6*^△^*^/^*^△^ cells. Neural lineage-inducing genes, such as *Sox1, Sox2*, and *Pax6,* were highly upregulated in *Ddx6*^△^*^/^*^△^ cells, whereas the mesendoderm lineage inducer *T* was significantly downregulated (Fig. 5F). We also conducted a monolayer differentiation experiment. ESCs favor neuronal differentiation in low-density, serum-free, and feeder-free culture conditions (Tropepe et al, 2001). Compared with WT ESCs, *Ddx6*^△^*^/^*^△^ ESCs differentiated and developed into neurons quickly. *Ddx6*^△^*^/^*^△^ cells exhibited stronger expression of TuJ1 with the morphology of well-developed dendrites and axons on differentiation Day1 (Fig. 5G). This observation was similar to premature neural differentiation noted in E8.5 *Ddx6*^△^*^/^*^△^ embryos.

Taken together, *Ddx6*^△^*^/^*^△^ embryos developed normally until the blastocyst stage and ESCs had no defects in self-renewal. However, the differentiation capacity of *Ddx6*^△^*^/^*^△^ pluripotent cells was strongly skewed to neuronal lineage commitment.

BMP signaling is important to prevent differentiation of ESCs to the neuronal lineage (Ying et al., 2003). Thus, strong inclination of *Ddx6*^△^*^/^*^△^ ESCs toward the neuronal cell fate would suggest that this brake is nonfunctional. We investigated whether SMAD-dependent BMP signaling pathway was repressed in *Ddx6*^△^*^/^*^△^ cells. In ESCs and the initial differentiation stage, BMP signaling has a transcriptional repressive role on these SMAD1/5 target genes (Fei et al., 2010). We examined expression pattern of known SMAD1/5 target genes in ESCs and found that the expression of *Accn4, Alx3, Dpysl2*, and *Kdm6b* was higher in *Ddx6*^△^*^/^*^△^ cells (Fig. 5H). These suggest that target genes were transcriptionally de-repressed due to reduced BMP signaling. In particular, *Dpsyl2* and the H3K27 demethylase *Kdm6b* are early neural differentiation regulators (Burgold et al., 2008; Fei et al., 2010); therefore, their high expressions are consistent with the phenotype of *Ddx6*^△^*^/^*^△^ ESCs preferring neural lineage commitment. We then asked whether aberrant transcriptional activation of a set of BMP signaling inhibitors also occurred in *Ddx6*^△^*^/^*^△^ pluripotent cells. Like *Ddx6*^△^*^/^*^△^ embryos, *Ddx6*^△^*^/^*^△^ ESCs exhibited a significant increase in expressions of negative regulators of the BMP pathway during EpiLC induction (Fig. 5I). Along with suppressed BMP signaling, *Ddx6*^△^*^/^*^△^ ESCs also displayed the features of enhanced Nodal signaling by having high expression level of *Nodal* and *Nanog* (Fig. 5D,E). These indicate that inhibition of BMP signaling and increase of Nodal signaling are the cell-intrinsic changes occurring when DDX6 is absent.

### Depletion of DDX6 quickly induce transcriptional upregulation of *Nodal* and the negative regulators of BMP signaling

To further confirm that the aberrant activation of BMP signaling inhibition is a cell-intrinsic property of *Ddx6*^△^*^/^*^△^ cells, we conditionally deleted *Ddx6* using the *Rosa-CreER^T2^; Ddx6^flox/flox^* mouse line. As we expected that loss of DDX6 to cause mesoderm formation defects, we removed DDX6 during gastrulation. Firstly, we examined the time required for complete depletion of DDX6. When tamoxifen was administered to the pregnant female via oral gavage at E6.5, *Ddx6* deletion and depletion of existing DDX6 proteins were completed by E7.5 (Fig. 6A). We then injected tamoxifen at E6.5 and collected embryos at E8.5 to examine their phenotypes. Conditional knockout (cKO) embryos of the same litter exhibited variable phenotypes like conventional knockout embryos (Fig. 6B). A few had marked posterior truncation (KO2, KO17). The others developed the mid-to-posterior body part, but it was shorter and smaller, and the head and heart morphology was abnormal (KO7, KO12). Although the phenotypes were milder than those of conventional KO embryos, conditional KO embryos also demonstrated characteristic gene expression of *Ddx6* mutants. The expression of the negative regulators of BMP (*Chrd, Noggin, Lgals9, Xdh, Pai*), *Nodal*, and *Eomes* increased after the depletion of DDX6 (Fig. 6C). Therefore, this conditional KO experiment reconfirmed that DDX6 is essential during gastrulation and that the loss of *Ddx6* makes cells activate inhibitory regulation of BMP signaling.

**Figure 6.**
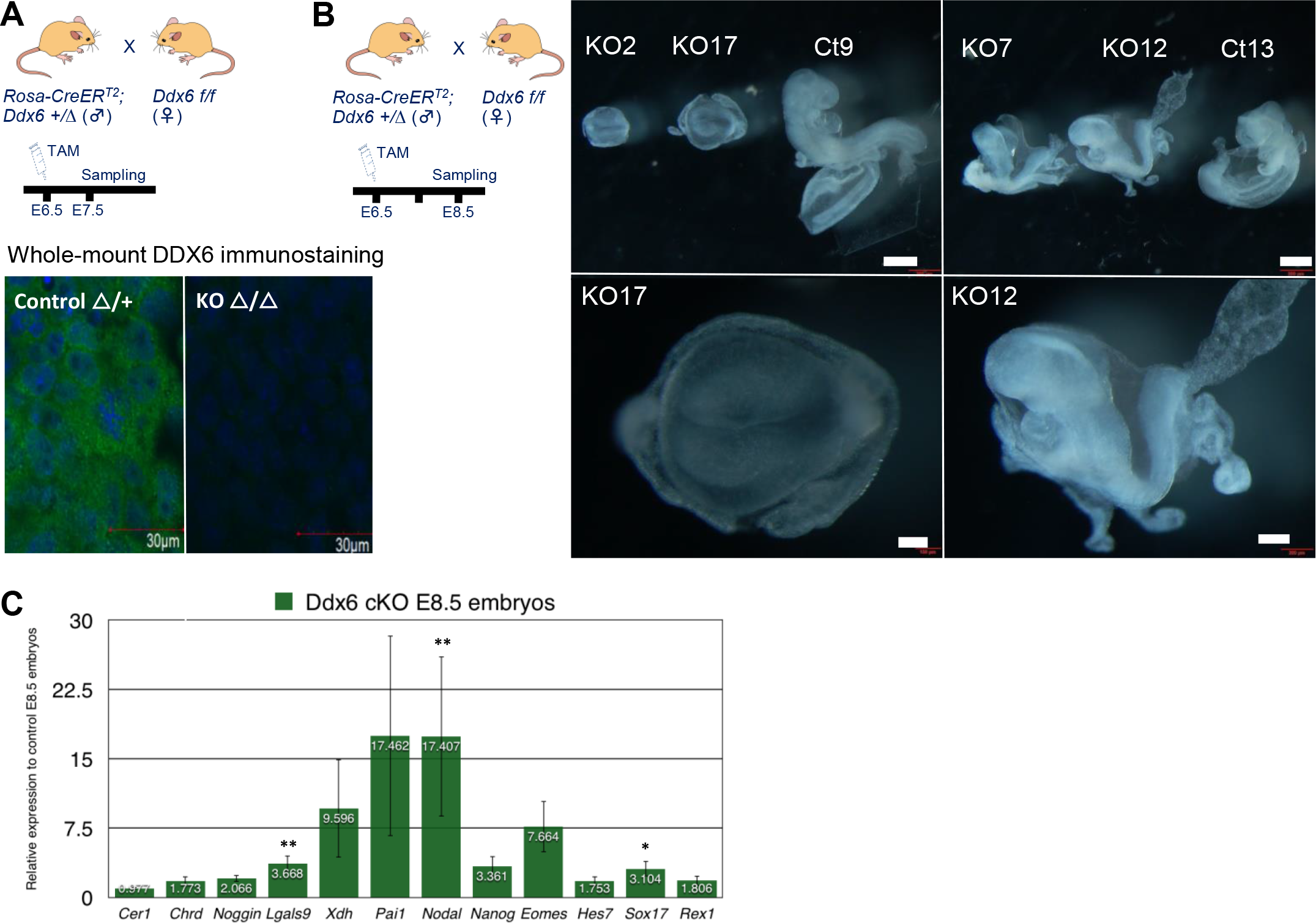
Conditional depletion of DDX6 quickly upregulates expressions of the BMP signaling inhibitors and *Nodal*. (A) Tamoxifen was injected at E6.5. Whole-mount DDX6 immunostaining confirmed that the complete depletion of DDX6 takes about 1 day (n = 3). (B) E8.5 cKO embryos exhibited similar phenotypes to conventional KO embryos (scale: 500 μm for group, 100 μm for KO17, 200 μm for KO12) (n = 6). (C) RT-qPCR analysis of several key genes in *Ddx6* cKO E8.5 embryos. The ratio of mutants with a milder phenotype was higher among cKO embryos, thus most embryos that were used for RT-qPCR analysis looked similar to the KO12 embryo. Mean ± SEM. The statistical significance was calculated by Wilcoxon rank sum test (n = 9∼10) (* *α* = 0.05 significance level, ** *α* = 0.01).

### Genetic dissection of the DDX6-mediated RNA regulatory pathways: DDX6 mainly works through the miRNA pathway during early embryogenesis

Next, we aimed to identify which DDX6 pathway is most crucial during early development. DDX6 functions as a hub of post-transcriptional regulation. Due to its wide range of involvement, it is difficult to pinpoint which pathway is responsible when DDX6 is depleted. Therefore, we individually disrupted three main DDX6-mediated pathways by knocking out a key gene of each pathway (Fig. 7A). Translational repression along with P-body formation was impaired in *Eif4enif1* KO, 5’-to-3’ mRNA degradation was impaired in *Dcp2* KO, and miRNA-mediated gene silencing was disrupted in *Dgcr8* KO (Andrei et al., 2005; Ferraiuolo et al., 2005; Wang et al., 2007; Aizer & Kalo, 2014; Ayache et al., 2015).

**Figure 7.**
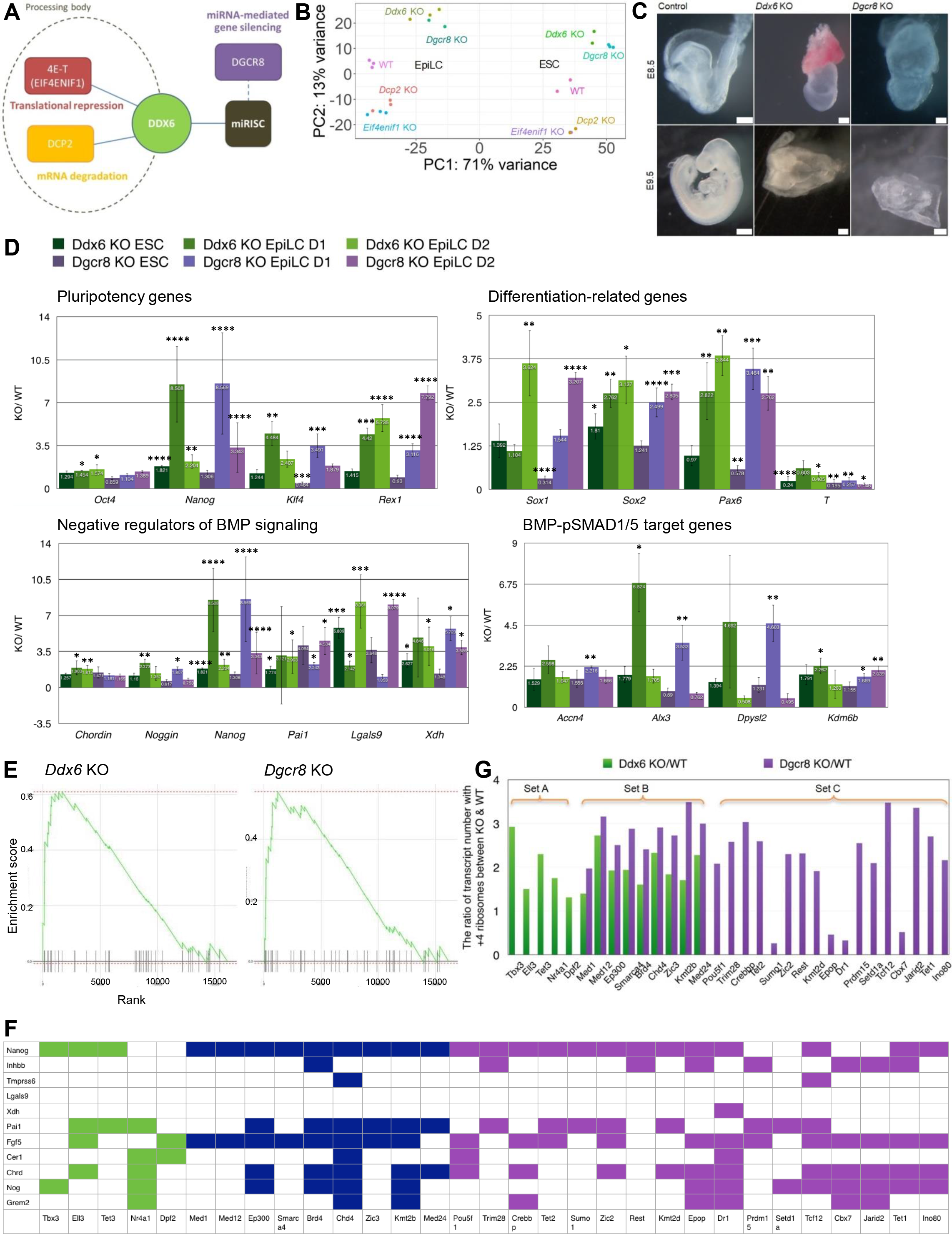
Genetic dissection of the DDX6 functions found that the DDX6-miRNA pathway has a crucial role during early embryogenesis. (A) A scheme of genetic dissection of the DDX6-mediated RNA regulatory pathways. Three major DDX6-mediated pathways were disrupted by knocking out the key gene of each pathway. (B) PCA plot of ESC and EpiLC Day2 of each genotype group. (C) *Dgcr8* KO embryos exhibit similar morphology to *Ddx6* KO embryos. (E8.5 images, scale: 200 μm for WT & *Ddx6* KO, 100 μm for *Dgcr8* KO, n ≥ 3; E9.5 images, 500 μm for WT, 200 μm for *Ddx6* KO & *Dgcr8* KO, n ≥ 3). (D) Comparison of gene expression between *Ddx6* KO and *Dgcr8* KO. RT-qPCR analysis of the expression trend of several key genes during EpiLC induction period. Each bar represents the relative expression of KO cells to WT cells at the indicated time point. Mean ± SEM. Student’s t test (n ≥ 3) (*p ≤ 0.05, **p ≤ 0.01, ***p ≤ 0.001, ****p ≤ 0.0001). (E) GSEA enrichment plot of the “negative regulation of BMP signaling pathway” gene set in *Ddx6* KO and *Dgcr8* KO ESC as compared to WT ESC. Black bars represent the position of the genes that are belong to this gene set (n = 45) in the whole ranked gene list. The green line shows the overall distribution of this gene set (whether over-represented at the top (left) or bottom (right) of the ranked list of genes). (F) The table showing which transcription factor (column) binds to which negative regulator of BMP signaling (row). Green color: Set A; Navy: Set B; Purple: Set C from Fig. 7G. (G) Analysis of the translation level change of transcription factors that bind to the BMP negative regulators. Only the transcripts that showed statistically significant change are shown in this graph (p ≤ 0.05). Set A was increased only in *Ddx6* KO (5 genes). Set B in both (9 genes) and Set C only in *Dgcr8* KO (17 genes).

To see the effects on the transcriptomic landscape over the time course of development, we generated cDNA libraries of ESC and EpiLC Day2 stage from all mutant groups and WT. PCA analysis demonstrated that *Ddx6* KO is greatly different from WT, *Eif4enif1* KO, and *Dcp2* KO, while it is very similar to *Dgcr8* KO (Fig. 7B). Moreover, *Dgcr8* KO embryos in a later developmental stage also showed very similar morphological defects to *Ddx6* KO embryos (Fig. 7C). *Eif4enif1* KO and *Dcp2* KO, whose protein functions directly affect P-body metabolism, resulted in similar transcriptomes. *Eif4enif1* KO, in which translational repression on transcripts is disrupted, resulted in disassembly of P-bodies, while *Dcp2* KO caused enlargement of P-bodies due to blockage of mRNA degradation (Fig. S4A). Even though their transcriptomes were different from WT, gastrulation occurred quite normally in KO embryos except subtle defects (Fig. S4B-C). E9.5 *Eif4enif1* KO embryos were smaller than the littermate control, and E9.5 *Dcp2* KO embryos exhibited smaller posterior body structure. Based on these data, we conclude that P-bodies are dispensable at least until the peri-gastrulation stage. Changes on P-body metabolism affected the transcriptome of cells, but the changes in gene expression were not substantial to alter the differentiation capacity of pluripotent stem cells.

Because of similar transcriptomic changes in cells and embryo phenotypes between *Ddx6* KO and *Dgcr8* KO, we conducted detailed analyses during ESC-to-EpiLC differentiation through RT-qPCR. *Dgcr8* KO cells exhibited very similar characteristics to *Ddx6* KO pluripotent cells: enhanced pluripotency and the stronger expression of the neural lineage-inducing factors with a decreased differentiation capacity to the mesendoderm lineage (Fig. 7D). As we have identified that repression of BMP signaling is the primary change of cellular condition in *Ddx6* KO, we checked if it also happens in *Dgcr8* KO. *Dgcr8* KO cells had upregulated expression of the negative regulators of BMP signaling and de-repression of pSMAD1/5 target genes.

Although GO term enrichment analysis of pluripotent cells RNA-seq did not hit ‘negative regulation of BMP signaling’ (Fig. S4D), a more sensitive gene set enrichment analysis (GSEA) detected this gene set. The ‘negative regulation of BMP signaling’ gene set was highly upregulated only in *Ddx6* KO and *Dgcr8* KO ESCs, but not in *Eif4enif1* and *Dcp2* KO ESCs (Fig. 7E, S4E). These indicate that DDX6-mediated RNA regulation and miRNA-mediated gene silencing share the common role, especially preventing aberrant transcriptional activation of the negative regulators of BMP signaling.

We next searched the underlying mechanism for increased transcript level of BMP inhibitors in *Ddx6* KO and *Dgcr8* KO. As DDX6 is suggested to participate in translational repression process mediated by a miRNA-induced silencing complex (miRISC) (Freimer et al., 2018), transcriptional upregulation of the negative regulators of BMP signaling would be the secondary effect. We hypothesized that translation of certain transcript factors is increased in the absence of translational repressive regulation by DDX6 and miRNAs, and eventually transcription of their target genes gets upregulated. To check this idea, we searched transcription factors that bind to the BMP negative regulators using ChIP-Atlas database (Oki & Ohta, 2015), and found some common proteins (Fig. 7F). Changes in translation were assessed by examining published *Ddx6* KO and *Dgcr8* KO ESC polysome profiling data (Freimer et al., 2018), and the number of transcripts with high polysome (+4 ribosomes), which are being translated, was increased in both KOs (Fig. 7G). Of note, most of them regulate *Nanog* transcription, and *Fgf5, Pai1, Chrd, Nog, Cer1, Grem2* are also frequently targeted by these transcription factors.

Therefore, this result supports our reasoning of the underlying mechanism that causes transcriptional upregulation of the negative regulators of BMP signaling in the absence of DDX6 and the miRNA-mediated gene regulation.

## Discussion

This study delineated an essential role of DDX6 in proper cell lineage specification and differentiation in early mouse embryogenesis (Fig. 8A-B). We found that DDX6 prevents cells from activating negative regulation of BMP signaling through the miRNA-mediated gene silencing. The genes that are related to ‘negative regulation of BMP signaling’ were upregulated in both *Ddx6* KO E8.5 embryos and ESCs. Their phenotypes including posterior body and mesoderm development defect, premature neural induction, and de-repression of BMP- SMAD1/5 target genes indicate that the BMP pathway is nonfunctional in *Ddx6* KO. Another characteristics of *Ddx6* KO was enhanced Nodal signaling, in an antagonistic relationship with BMP signaling. Promoted pluripotency, such as increased *Nanog* expression, can be attributable to high *Nodal* expression.

**Figure 8.**
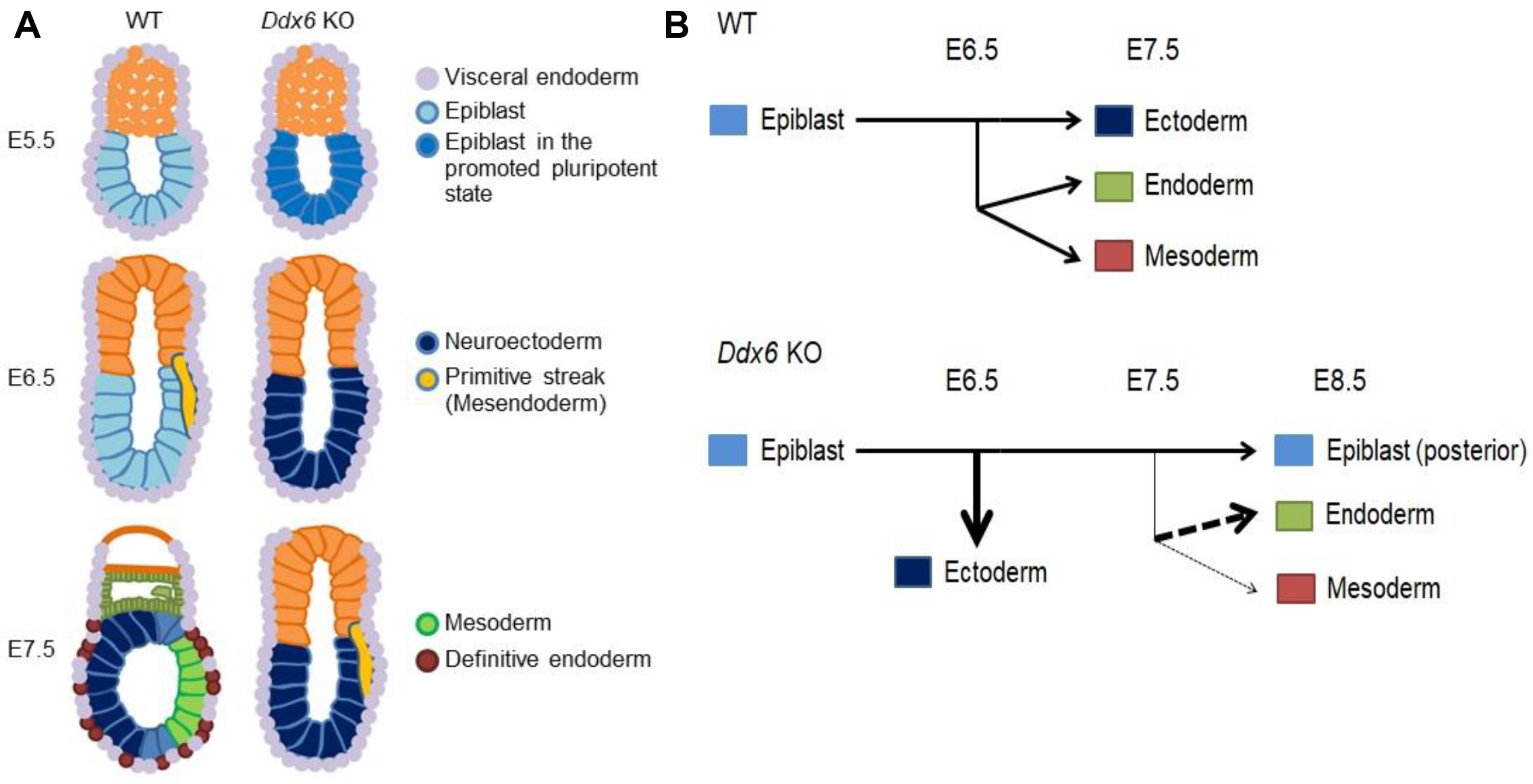
Schemes describing developmental defects caused by loss of DDX6-mediated RNA regulation. (A) Development of the three primary embryonic germ layers is largely affected by DDX6. Neuroectoderm is specified earlier than WT whereas the formation of primitive streak is delayed (The smaller size of *Ddx6* mutant is not reflected on the images). (B) Changes in cell-lineage specification from pluripotent stem cells caused by *Ddx6* loss is depicted on a horizontal diagram. Uncommitted *Ddx6*^△^*^/^*^△^ pluripotent cells posses promoted pluripotency and strongly favor commitment to the neuronal lineage. In WT embryos, mesendoderm lineage arises at ∼E6.5 as the primitive streak is formed, and three germ layers are simultaneously developed at ∼E7.5. In *Ddx6* KO embryos, premature neural induction occurs while one day delay of the primitive streak formation is observed. During mesendoderm segregation, definitive endoderm specification is increased whereas mesoderm specification is greatly reduced due to the patterning defect of the primitive streak. Posterior epiblast cells cannot exit the pluripotency on time, which would also impede differentiation processes.

Single cell multi-omics analysis of peri-gastrulating mouse embryos revealed that enhancers of the ectoderm lineage are active in the epiblast of E4.5 embryos (Argelaguet et al., 2019), but the mechanism that enables ectoderm specification to occur at the same time as the mesoderm and endoderm develop during gastrulation remains unclear. Genome-wide mapping of SMAD1/5 targets revealed that BMP-SMAD pathway mainly represses transcription in mESCs (Fei et al., 2010). SMAD1/5 binds to the promoter of a set of developmental regulators whose expression affects cell fate determination. Among them, those related to nervous system development were most significantly enriched. Taken together with our observations of *Ddx6* KO, in which BMP signaling is repressed, transcriptional repression by the BMP-SMAD1/5 pathway may be the main mechanism that prevents precocious activation of neuroectoderm (NE) genes in the open chromatin state and allows simultaneous development of the three germ layers. We also contemplated about the relationship between pluripotency and neural lineage. Of note, the deletion of *Ddx6* promoted both pluripotency and inclination to neural lineage commitment. This serves as further evidence that the pluripotent state and cell-fate decision to neuroectoderm are somehow managed together. In addition, we found that the well-known NE or neuronal marker TuJ1 (TUBB3) is expressed in the primed pluripotent epiblast (Fig. S5A). This suggests the closeness of pluripotent cells and neuroectoderm-committed cells. Further understanding of this property will be useful for stem cell and neurodevelopmental biology research.

As embryos have abundant P-bodies from blastocyst to peri-gastrula embryos, the marginal effect of disrupted P-body metabolism on early embryogenesis was unexpected. Di Stefano et al. (2019), stated that disruption of P-bodies is the reason of *Ddx6*-deficient ESCs being “hyper-pluripotent,” but we refute this idea based on our results. Deletion of *Eif4enif1*, one of the core P-body component, led to disassembly of P-bodies in ESCs, but *Eif4enif1* KO pluripotent cells did not display “hyper-pluripotent” property unlike *Ddx6* KO cells, and *Eif4enif1* KO embryos did not show developmental defects that *Ddx6* mutants displayed.

Therefore, P-bodies are not essential for development at least until the gastrulation stage. Di Stefano et al., knocked out *Lsm14a,* another core P-body component and a translational repressor, to examine whether *Ddx6* KO phenotype was P-body-dependent. Thus, it is possible that some specific functions of the DDX6-LSm14A complex are important rather than formation of P-bodies.

Our transcriptomic analyses identified the similarity between *Ddx6* KO and *Dgcr8*KO, but not with *Eif4enif1* KO or *Dcp2* KO. Only *Dgcr8* KO closely phenocopied *Ddx6* KO, indicating that DDX6 mainly works through the miRNA pathway among its various regulatory means during early embryogenesis. In *Dgcr8* KO, the miRNA-mediated gene silencing becomes nonfunctional because of failure of generating miRNAs (Wang et al., 2007). DDX6 participates in the effector step of miRNA-mediated gene silencing (Chen et al., 2014). Freimer et al. (2018), showed that loss of DDX6 only impairs miRNA-induced translational repression but not mRNA destabilization in mESCs. Thus, we have analyzed translation change of transcription factors that bind to the BMP negative regulators and found that their translation level was increased. They were categorized into three groups (Fig. 7G): Set A (only up in *Ddx6* KO), Set B (commonly up), Set C (only up in *Dgcr8* KO). Interestingly, there was much greater number of transcripts whose translation was increased only in *Dgcr8* KO (Set C). It has been thought that DDX6 would involve in all translational repression activities mediated by miRISCs, because its interaction with miRISCs occurs through the CCR4-NOT complex whose binding to miRISCs is essential for their actions (Braun et al., 2011; Fabian et al., 2011; Zekri et al., 2013; Chen et al., 2014). However, the clear segregation of three classes would imply that DDX6 has specificity of translational repression on the certain transcripts rather than being a default component of miRISCs. Therefore, our analyses supplement the mode of DDX6 action in miRISC-mediated translational repression. This study also signifies the strong impact of the miRNA-mediated gene regulation on transcription factors and signaling pathways during early embryogenesis.

Lastly, RNA-seq of E8.5 *Ddx6*^△/△^ mouse embryos revealed the similar features to *Ddx6*-deficient human somatic cells. Lumb et al. (2017), reported that DDX6 is required to prevent the aberrant activation of interferon-stimulated genes (ISGs). Many immune response-related terms were enriched in the GO analysis of highly upregulated genes in *Ddx6* mutant embryos (Fig. 2A). In addition, ISG genes, such as *Ifit1*, *Ifitm1,* and *Oas1,* were upregulated in *Ddx6*^△/△^ embryos, as in the above report (Fig. S5B). Thus, our study reinforces that DDX6 plays an important role in regulating immune response-related genes. We provide a further detailed observation. When comparing RNA-seq data of *Ddx6* KO and *Dgcr8* KO ESCs, immune response-related GO terms were enriched in the genes that were commonly upregulated in *Ddx6* KO and *Dgcr8* KO or only in *Dgcr8* KO. This suggests that miRNAs have an important function regulating immune gene expression, and DDX6 again works together with the miRNA system to suppress immune gene activation. Another notable similarity is the function of DDX6 as an inhibitor of aberrant activation of a certain set of genes. Cells lacking DDX6 activated ISG expression in the absence of external interferon stimulation; in our case, *Ddx6*^△/△^ cells activated expression of negative regulators of BMP signaling to make themselves less responsive to external BMP stimulation. Cells have the ability to sense and quickly react to their environments. However, how they coordinate the intrinsic gene regulatory network with extrinsic stimuli remains elusive. Based on these observations, we conjecture that DDX6-mediated post-transcriptional RNA regulation may become an important link between intracellular processes and extracellular stimuli. This feature also emphasizes the important role of DDX6 as a gene expression regulator. The studies on DDX6 have wide applicability because it is expressed in various cell types and different developmental stages. Here we identified its embryogenic role and clearly segregated its molecular functions through a genetic dissection approach. Having insights on various DDX6 functions and corresponding molecular mechanisms would make this general RNA helicase a valuable target of gene regulation.

## Materials and Methods

### Mice

Mice were housed in a specific-pathogen-free animal care facility at the National Institute of Genetics (NIG). All experiments were approved by the NIG Institutional Animal Care and Use committee. *Ddx6*^△^*^/+^*, *Ddx6^flox/flox^*, *Rosa-CreER^T2^*, *Ddx6-mCherry, Eif4enif1* KO*, Dgcr8* KO*, Dcp2* KO mice were used in this study. The production strategy of *Ddx6^flox/flox^* mouse line is described in (Kato et al., 2019). *Ddx6*^△^*^/+^* mouse line was generated by electroporation of two gRNAs (targeting AACAAAGCCAACCCGGGACA and CTATGTGCTGTAGCTTAGTC) and Cas9 protein into fertilized eggs resulting in the removal of exon2. Genotyping was done by primers [Ddx6-LA-Fw1: TTGTGCTGGGATGAGCCTAC; Ddx6-RA-Rv1: AGTTGCATCAACGACAGGAGAG]. *Ddx6-mCherry* reporter mouse was established in NIG by injecting a targeting vector containing mCherry with homology arms of *Ddx6* gene with Cas9-gRNA designed at the C-terminal of *Ddx6* gene and Cas9 protein. Genotyping was done by primers [mCherry-L1: GGAACAGTACGAACGCGCCG; DDX6-GR1: GACAGGTGCATGTGTTCACCC]. *Eif4enif1* KO mice and *Dgcr8* KO mice were directly obtained as F0 generation, which were produced by delivering Cas9 protein and guide RNAs targeting (“TCTGGTTCATACCGTAGTTT”, “AACTTACTTTCGTATAGCGA” for Eif4enif1 exon2) and (“TGAATCCTAATTGCACCCGT”, “GAACAGGAAGCATACGGGTA”, “TGGGTCGGTCTGCAGAGTTG” for *Dgcr8* exon4 & 5) into the fertilized eggs via electroporation. Similar strategy was used to establish *Dcp2* KO mouse line, two gRNAs targeting “AACAAAGCCAACCCGG” and “CGCGGCACTGAAGTGT” and Cas9 protein were delivered via electroporation. The mouse line that has correctly deleted exon2 was selected and expanded. The homozygous KO pups were acquired by crossing *Dcp2* heterozygous mice.

• For conditional deletion of the floxed *Ddx6* alleles, 600 μL of 10 mg/mL tamoxifen was administered to the pregnant females.

### Establishment of ES cells

• *Ddx6* KO: *Ddx6*^△^*^/+^* mice were intercrossed and blastocysts were collected from the uterus on E3.5. Collected blastocysts were cultured on mitomycin-treated mouse embryonic fibroblast feeder cells in 2i-LIF medium (ESGRO Complete Basal medium (Millipore, Germany) supplemented with leukemia inhibitory factor (Wako, Tokyo, Japan), 0.4 μM MEK inhibitor PD0325901 (Wako, Tokyo, Japan), 3 μM GSK3 inhibitor CHIR99021 (Wako, Tokyo, Japan) and Penicillin-Streptomycin (Invitrogen). Blastocyst outgrowths were disaggregated and passaged onto the new wells plated with feeder cells in the same medium condition. Once ES cell colonies developed, they were expanded for genotyping and the storage. Genotyping was done with primers [Ddx6-LA-Fw1: TTGTGCTGGGATGAGCCTAC; Ddx6-RA-Rv1: AGTTGCATCAACGACAGGAGAG].

• *Dcp2* KO: Firstly, *Dcp2* cKO ES line was established by replacing the endogenous exon2 with the floxed exon2 through CRISPR/Cas9-mediated homologous recombination. *Dcp2* KO ES line was acquired by incubating cKO ESCs in culture medium containing 4-Hydroxytamoxifen (4- OHT). Genotyping was done with primers [Dcp2-LA-Fw1: TTCTGCTGCTTTCAAGCCTGG; Dcp2-int2-R2: ACATTCGCTACAACAACGCTTC].

• *Eif4enif1* KO: was also generated by deleting floxed exon2 from conditional KO ESCs, established by replacing the endogenous exon2 with the floxed exon2 through CRISPR/Cas9- mediated homologous recombination. Genotyping was done with primers [4ET-int1-F1: GTGACAGGCACTTTCCAGCAG; 4ET-int2-R1: TTCCAAAGCCTTAGCTGCTTCTC].

• *Dgcr8* KO: was established by deleting exon4 and a part of exon5 using two Cas9 vectors targeting “TGAATCCTAATTGCACCCGT” and “TGGGTCGGTCTGCAGAGTTG.” CRISPR direct (http://crispr.dbcls.jp/) was used to find Cas9 target sites, and the target sequence was integrated into a modified px330 Cas9 vector (addgene), which contains the pgk-puromycin cassette. ESC transfection was done with Lipofectamine 2000 (Invitrogen). Genotyping was done with primers [Dgcr8-int3-F2: GCTCCTGGAGTAGGCATGTTG ; Dgcr8-ex5-R1: TTCACTTGTCCCAGGGCTCC].

### Immunostaining

• For frozen section immunohistochemistry, embryos were fixed in 4% paraformaldehyde for 30 min at 4°C, submerged in 10% sucrose for 1∼2 hours, in 20% sucrose overnight at 4°C, and frozen in Tissue-Tek O.C.T. compound (Sakura Finetek, Tokyo, Japan). Each 6 μm-thickness section was applied to glass slides. After blocking with 3% skim milk in PBS-T (PBS with 0.1% Tween20 (Sigma-Aldrich)) at room temperature for 45 min, samples were incubated with primary antibodies overnight at 4°C. The primary antibodies are listed in Table 1. Next day, samples were washed with PBS-T and incubated with secondary antibodies labeled with Alexa Fluor 488, 594, 647 (1:1000 dilution in PBS-T, Invitrogen) for 1 hr 10 min at room temperature. DNA was counterstained with DAPI (100 ng/mL). Images were acquired by the Olympus FV1200 confocal microscope and processed with FV10-ASW (version 4.0) software.

• For whole-mount immunostaining, embryos were fixed in 4% paraformaldehyde for 30 min at 4°C, and permeabilized with 1% Triton X-100 in PBS for 30 min. Blocking was done with 10% FBS and 1% BSA for 1hr at room temperature. Samples were then incubated with primary antibodies for one overnight at 4°C, and secondary antibody reaction was done for another overnight at 4°C. Images were taken by the Olympus FV1200 confocal microscope.

• For immunocytochemistry, cells were fixed in 4% paraformaldehyde for 12 min at room temperature, then permeabilized with 0.3% Triton X-100 in PBS for 12 min at room temperature and blocked with 3% skim milk in PBS-T for 45 min at room temperature. After blocking, cells were incubated with primary antibodies overnight at 4°C. Next day, samples were incubated with secondary antibodies for 1 hr at room temperature and counterstained with DAPI. Images were acquired by the Olympus FV1200 confocal microscope or Leica DM6000 FS light microscope.

### Whole-mount *in situ* hybridization

Probe generation and whole-mount embryo ISH procedures were followed by the protocol described before (Biris and Yamaguchi, 2014).

### RNA-seq

• E8.5 embryos RNAs were collected from E8.5 embryos using TRIzol Reagent (Thermo Fisher Scientific). Genotyping was done with extraembryonic tissue or yolk sac and further confirmed by qPCR after acquiring cDNA. 300 ∼ 420 ng of RNA was used for E8.5 cDNA library generation with the KAPA-Stranded mRNA-seq kit (Illumina Platforms, KR0960-v5.17).

• ESC & EpiLCs of WT, *Ddx6* KO, *Eif4enif1* KO, *Dcp2* KO, and *Dgcr8* KO RNA from ESC and EpiLC Day2 samples were extracted using RNAiso Plus (Takara, Tokyo, Japan) according to manufacturer’s instruction. RNAs were selected by polyA. Two ESC cDNA libraries (three *Dgcr8* KO ESC) & three EpiLC Day2 cDNA libraries were generated for each genotype group. For *Dgcr8* KO EpiLC case, the reads of one library suggested possible contamination, so we excluded that library. n = 2 for *Dgcr8* KO EpiLC. cDNA libraries were generated using TruSeq Stranded mRNA (illumina, 20020595) following the accompanied protocol. Samples were sequenced by NovaSeq 6000 (101 bp paired-end sequencing). Average 47 million read pairs per sample.

### Bioinformatics analysis

• E8.5 embryos For all libraries, Low-quality sequences, adapters and polyA or T were trimmed or removed using Cutadapt (version 2.8) (Martin, 2011) with the following options: “-e 0.1 -q 20 -m 20 -O 3 -a GATCGGAAGAGCACACGTCTGAACTCCAGTCAC -a A{100} -a T{100}”. The raw reads and processed reads were checked using FastQC (version 0.11.7, http://www.bioinformatics.babraham.ac.uk/projects/fastqc/). To map the reads to the mouse reference genome, the UCSC mm10 mouse reference genome (fasta) and gene annotation (General Transfer Format (GTF)) file were downloaded from Illumina iGenomes (https://sapac.support.illumina.com/sequencing/sequencing_software/iGenome.html). To increase the mapping accuracy of splicing reads, splicing-site and exon information were extracted from the gene annotation GTF file using the Python scripts hisat2_extract_splice_sites.py and hisat2_extract_exons.py, respectively, from the HISAT2 (version 2.1.0) package (Kim, 2015). The HISAT2 index files of the reference genome were built including the extracted genomic information using “hisat2-build” command with options: “--ss” and “--exon”. Clean reads were then mapped to the HISAT2 index files using the HISAT2 with default options. The obtained Sequence Alignment Map (SAM) files were sorted by genomic coordinates and converted to Binary Alignment Map (BAM) files using SAMtools (version 1.9) (Li et al., 2009) “sort” command with option: “-O BAM”. Raw read counts per gene were calculated using featureCounts (version 2.0.0) (Liao et al., 2014) with options: “-s 2 -t exon -g gene_id -a iGenomes/mm10/Annotation/Genes/genes.gtf”. Normalized counts were calculated by the trimmed mean of M-values (TMM) method using edgeR (version 3.28.1) (Robinson et al., 2009) on R (version 3.6.3) [https://cran.r-project.org/]. Principal component analysis (PCA) were performed on the log2 transformed normalized counts obtained from the “cpm” function in the edgeR using the “prcomp” function in the R with default options. The detection of differentially expressed genes (DEGs) were performed using the edgeR with the cut-off criteria of log2 (fold change) > 1 or < -1. Gene ontology term enrichment analysis was done via Metascape (Zhou et al., 2019).

• ESC & EpiLC of WT, *Ddx6* KO, *Eif4enif1* KO, *Dcp2* KO, and *Dgcr8* KO

For all libraries, Low-quality sequences, adapters were trimmed or removed using fastp (version 0.20.0) (Chen et al., 2018) with the following options: “-G -3 -n 1 -l 80”. For preparation for mapping reads to the mouse reference genome, the Ensembl mouse reference genome (release- 102, Mus_musculus.GRCm38.dna_sm.primary_assembly.fa.gz) and the gene annotation GTF file (Mus_musculus.GRCm38.102.gtf.gz) were downloaded from the Ensembl ftp site (http://ftp.ensembl.org/). To increase mapping accuracy of splicing reads, splicing-site and exon information were extracted from the gene annotation GTF file using the Python scripts hisat2_extract_splice_sites.py and hisat2_extract_exons.py, respectively, from the HISAT2 (version 2.2.1) package. The HISAT2 index files of the reference genome were built including the extracted genomic information using “hisat2-build” command with options: “--ss” and “-- exon”. Clean reads were then mapped to the HISAT2 index files using the HISAT2 with default options. The obtained SAM files were sorted by genomic coordinates and converted to BAM files using SAMtools (version 1.13) “sort” command with option: “-O BAM”. Raw read counts per gene were calculated using featureCounts (version 2.0.3) with options: “-s 2 -p -- countReadPairs -B --primary -t exon -g gene_id -a Mus_musculus.GRCm38.102.gtf”, and then low-abundance genes were removed by removing the genes with total number of mapped reads < 10 among 26 samples. Normalized counts and statistical values for differential gene expression analysis were calculated by the default settings through the steps: 1. estimation of size factors, 2. estimation of dispersion, and 3. Negative Binomial GLM fitting and Wald statistics using the Bioconductor DESeq2 packages (version 1.32.0) (Love et al., 2014) in the R (version 4.1.1). For checking the gene expression correlation between samples, the pair-wise scatter plot was produced using log2(the normalized counts + 1), the “cor” function with the parameter “method=’spearman’, use=’pairwise.complete.obs’” and ggplot2 packages (Wickham, 2016) in the R. For PCA, the variance stabilizing transformed (vst) normalized counts were calculated using the vst function of the DESeq2 with the default settings and PCA was performed with the top 500 most variable genes using the DESeq2’s plotPCA function. For checking DEGs, MA plots for each comparison between sample groups were produced using the results of the DESeq2 analysis and the ggplot2. The detection of DEGs were performed using the DESeq2 with the cut-off criteria of adjusted p-value < 0.05 and log2(Fold Change) > 2 or < -2. Gene ontology enrichment term analysis was done via Metascape.

### Gene set enrichment analysis

Gene set enrichment analysis was implemented via the R package ‘fgsea’ (Subramanian et al., 2005; Sergushichey, 2016) using the genes pre-ranked on the basis of Wald statistic values of ESCs obtained from DESeq2 as ‘stat’. Used gene set was the ‘Biological Processes’ gene set collection from MSigDB v7.4 (Liberzon et al., 2011).

### Polysome profiling data analysis

Polysome profiling RNA-seq data of *Ddx6* KO and *Dgcr8* KO ESCs were obtained from Freimer et al., 2018 (GSE112767). Translation level was assessed by examining high polysome (4+ ribosomes) counts. Statistical significance was tested by student’s t-test.

### RT-qPCR

Embryos were frozen in RNAiso Plus (Takara) and extracted according to manufacturer’s instruction. RNA of cultured cells was extracted by RNeasy Mini Kits (Qiagen, Germany). Extracted RNA was treated with Recombinant DNaseI (Thermo Scientific) for 30 min at 37°C, and processed for reverse transcription using SuperScript III or IV Reverse Transcriptase (Invitrogen). Quantitative PCR was performed using KAPA SYBR Fast qPCR Kits (Nippon Genetics, Japan) on Dice Real Time System Single Thermal Cycler (Takara) or CFX96 Real-Time System (BioRad) machine. The primer sequences are listed in Table 2. The expression level was normalized to *Gapdh* gene and the relative expression was calculated by ΔΔC_T_ method.

**Table 2.**
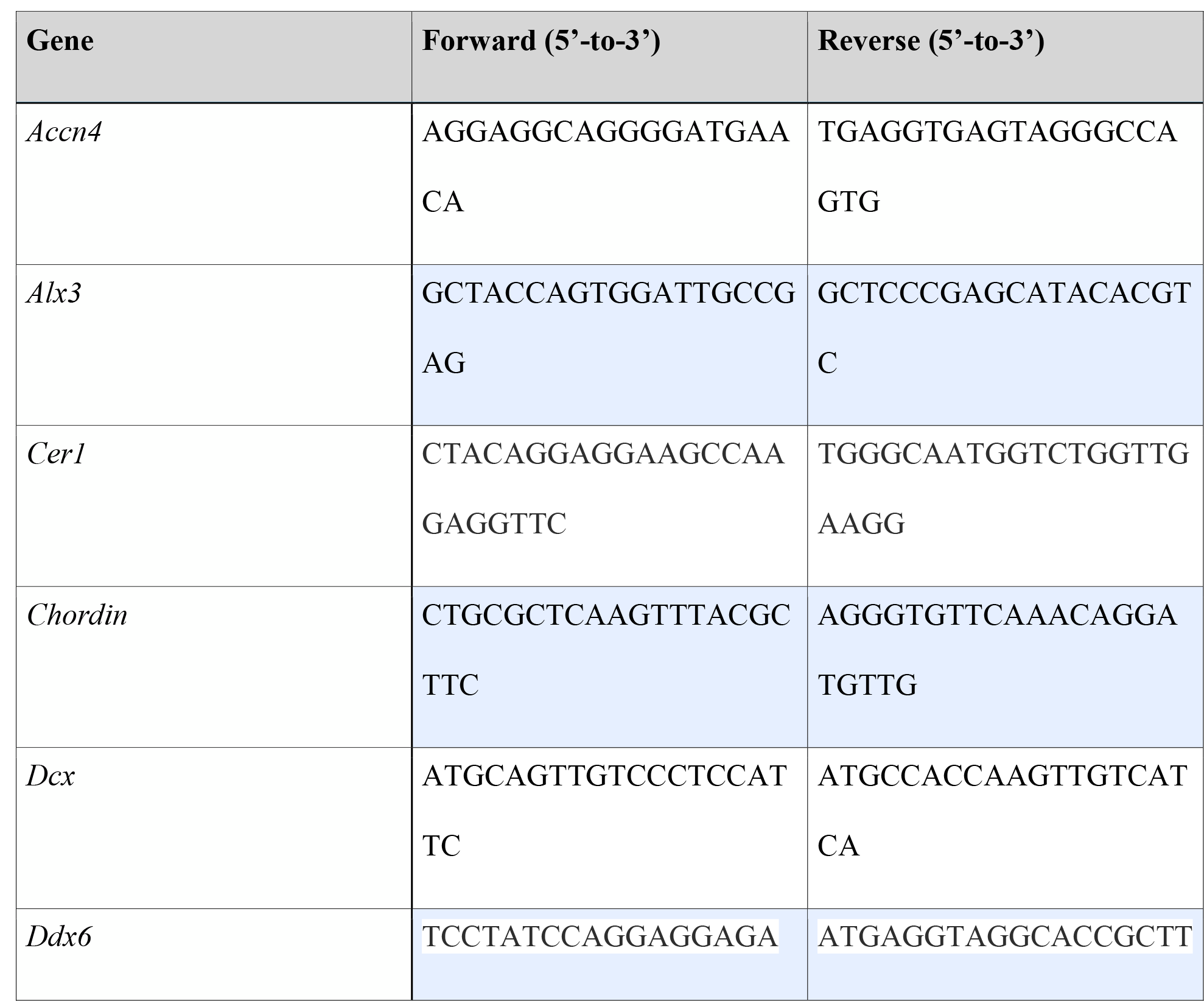
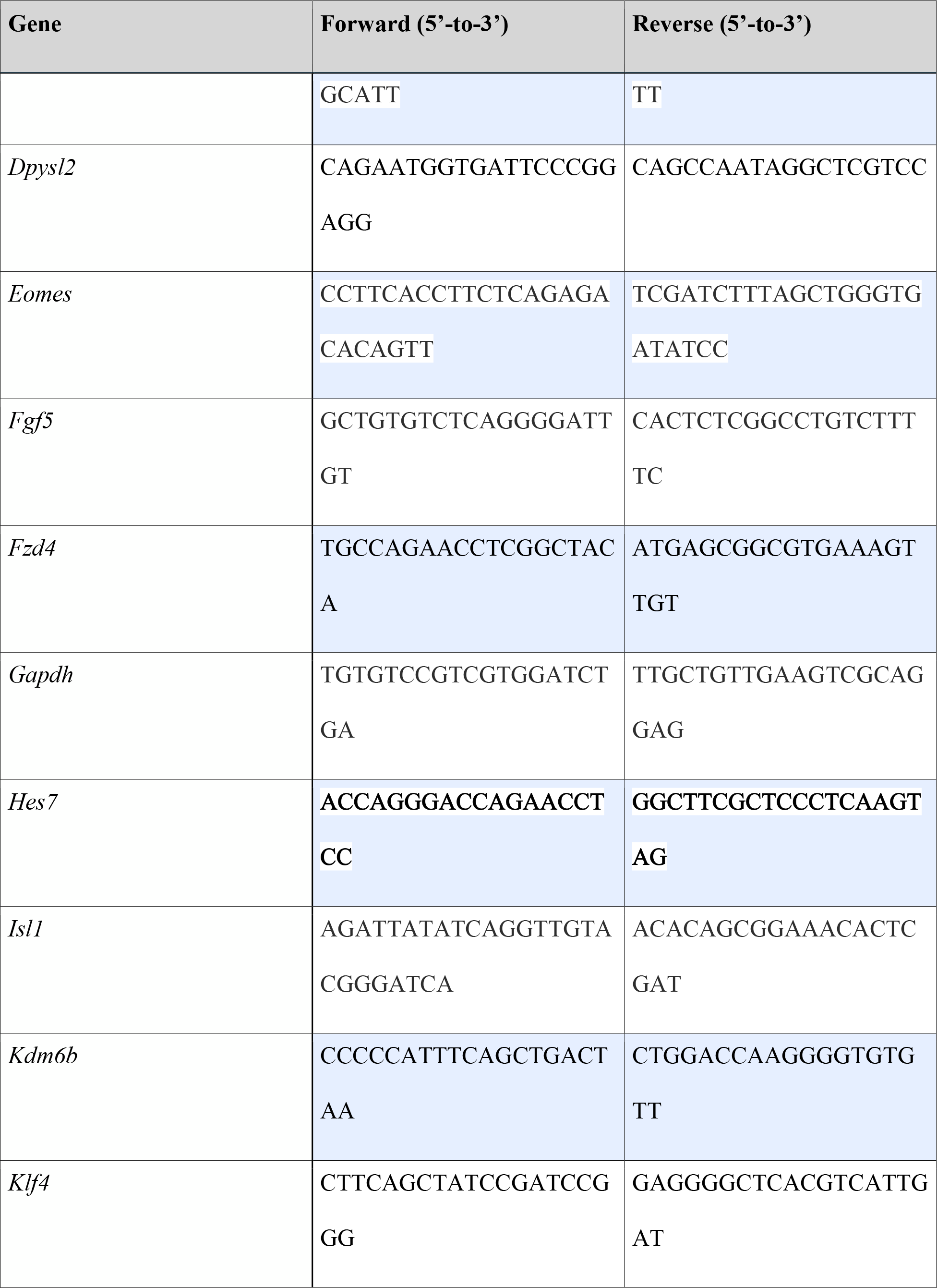
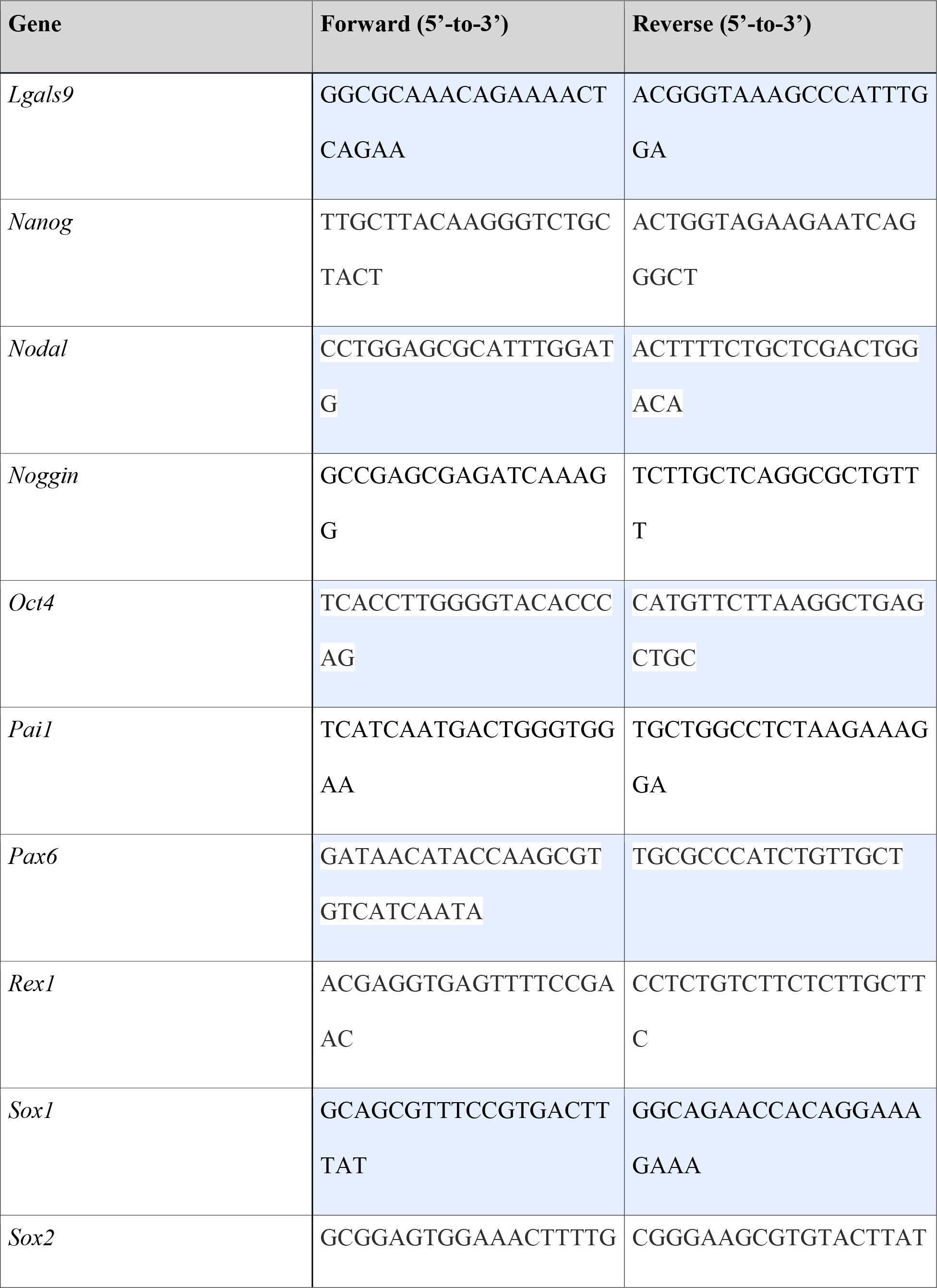
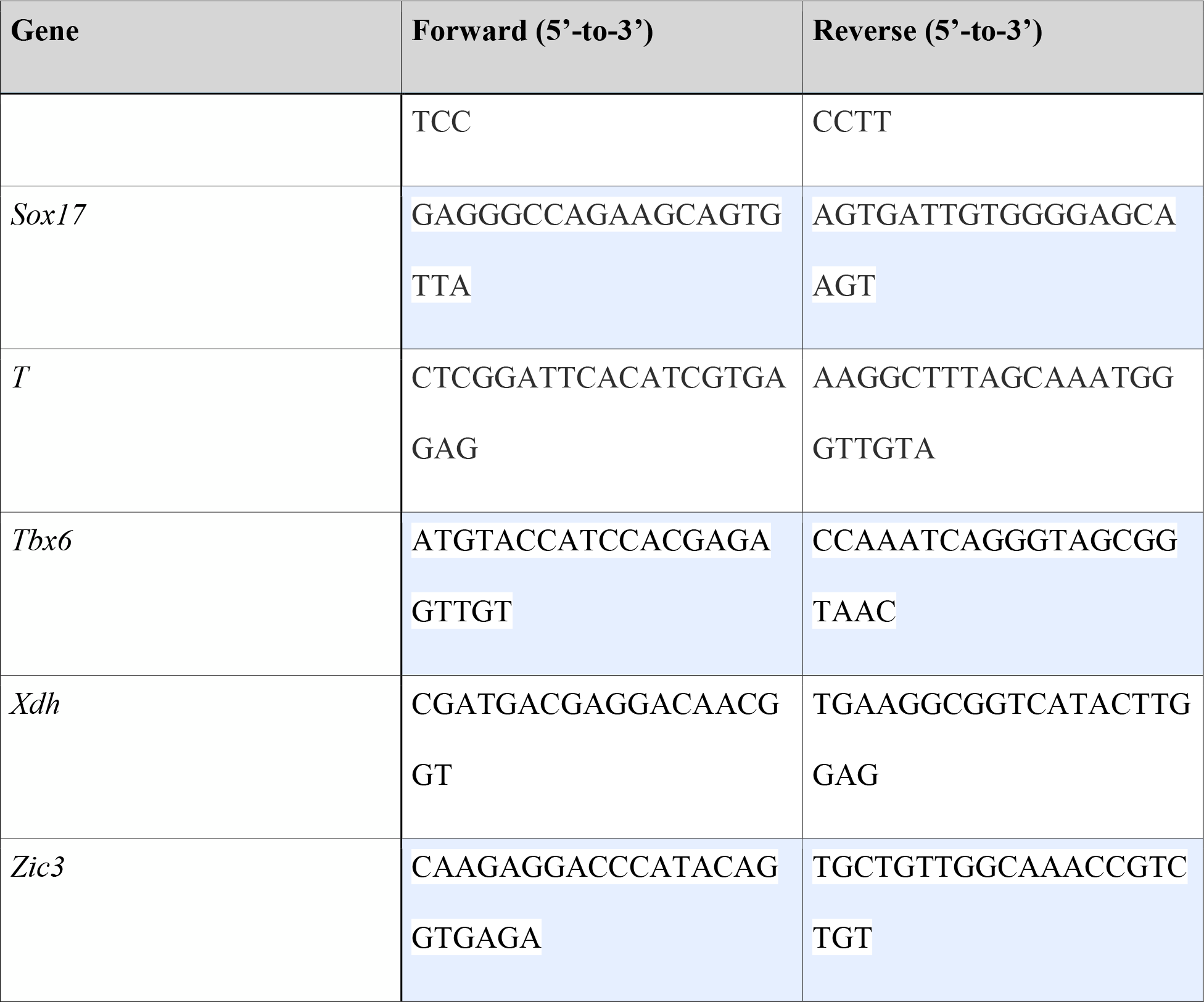
qPCR primer list.

### Statistical analysis

Statistical significance for *in vitro* experiments was examined by the Student’s t-test. *p ≤ 0.05, **p ≤ 0.01, ***p ≤ 0.001 and ****p ≤ 0.0001. Statistical significance for embryo RT-qPCR experiments was assayed by Wilcoxon rank sum test. *α = 0.05, **α = 0.01. Error bars represent s.e.m.

### ESC-to-EpiLC induction

A detailed protocol is described in (Hayashi et al., 2011). After feeder depletion, 1.3 x 10^5^ ESCs were plated on the well of 12-well plate pre-coated with fibronectin for 1 hr at 37°C. Medium was changed every day.

### Monolayer differentiation

After feeder depletion, 7 x 10^4^ ESCs were plated on the well of 24-well plate pre-coated with gelatin. Cells were incubated with ESGRO Complete Basal medium (Millipore, Germany).

### Data availability

• E8.5 embryo RNA-seq data have been deposited in the Gene Expression Omnibus (GEO) under accession code GSE171156.

• ESC & EpiLC RNA-seq data have been deposited in the Gene Expression Omnibus (GEO) under accession code GSE187390.

## Acknowledgments

We appreciate Prof. Ken Kurokawa for the support of transcriptomic analyses work. We also thank Dr. Danelle Wright for proofreading and Ms. Kumiko Inoue for supporting the mouse care.

## Supplementary figure legend

**Supplementary Figure 1.**
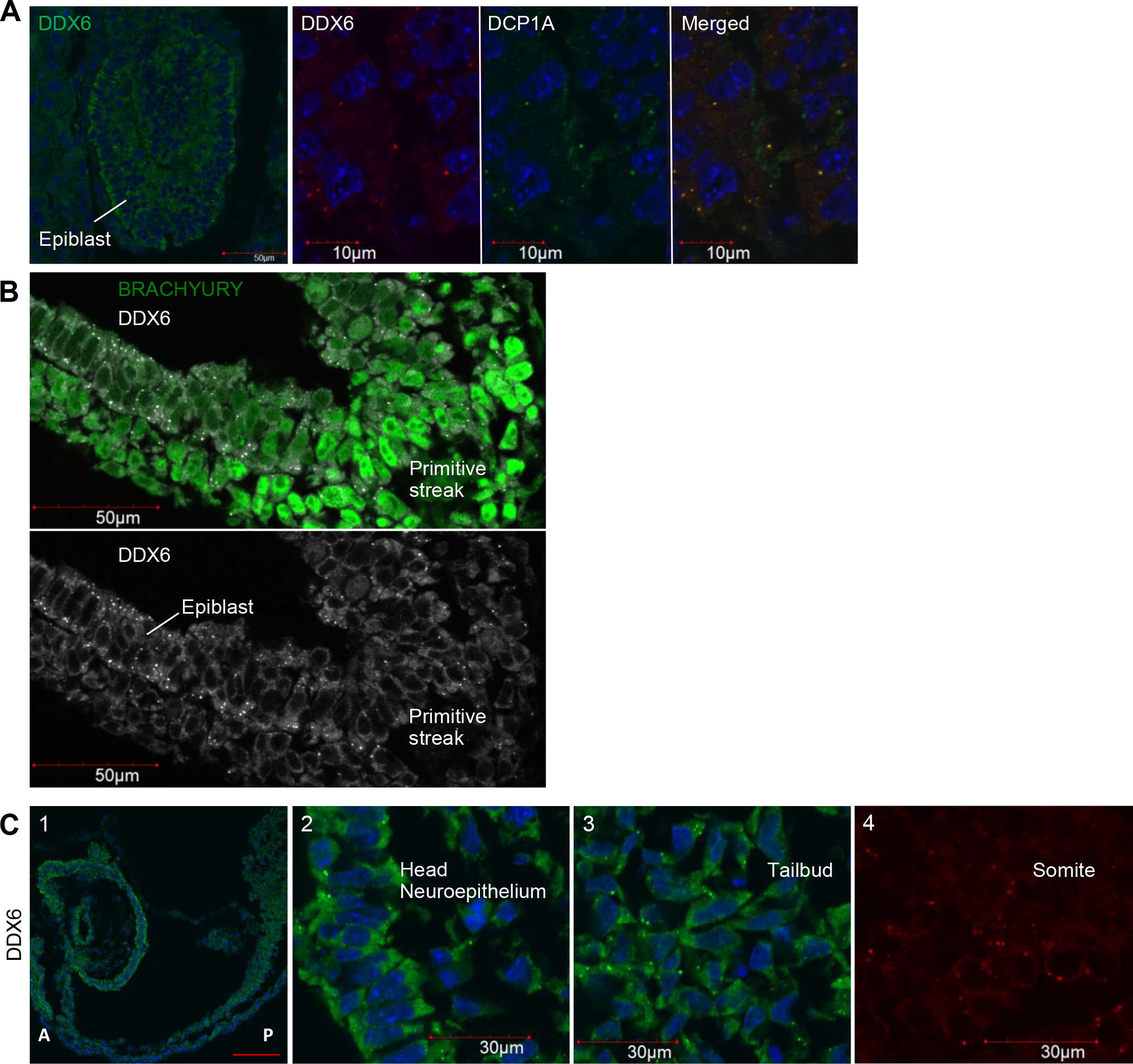
DDX6 expression in early embryos. (A) E6.5 embryo frozen section IHC for DDX6 and DCP1A. (B) E7.5 embryo frozen section IHC for BRACHYURY & DDX6. (C) (1-3) E8.5 embryo frozen section IHC for DDX6. (4) Imaging of E8.5 embryo expressing DDX6-mCherry (scale for (1): 100 μm).

**Supplementary Figure 2.**
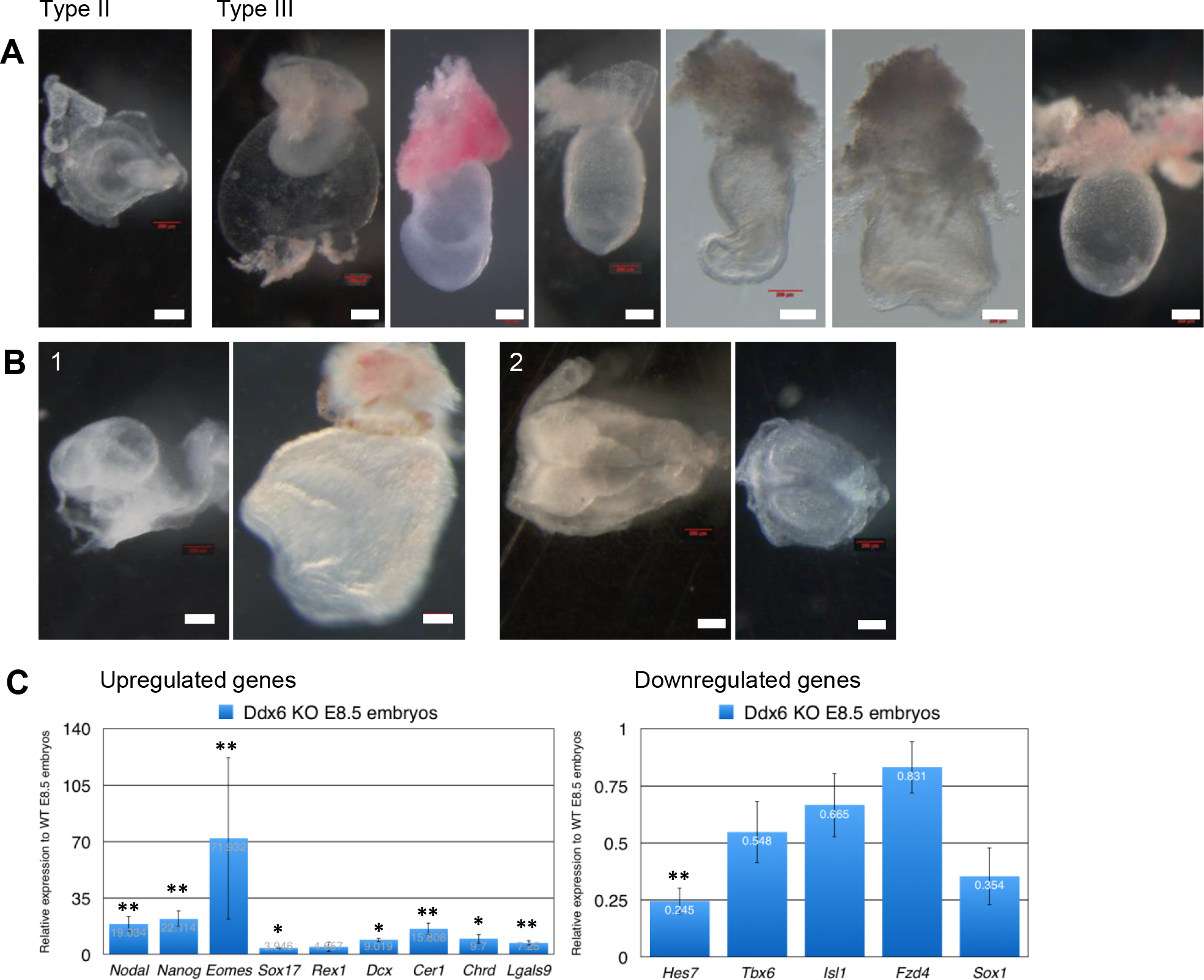
Detailed phenotypes of *Ddx6* KO embryos. (A) Variations in the morphology of E8.5 *Ddx6* KO embryos (scale: 200 μm). (B) Two types of E9.5 *Ddx6* KO embryos: (1) some mid-posterior body developed, (2) severe posterior truncation (scale: 200, 100, 200, 200 μm). (C) RT-qPCR analysis of some key genes in *Ddx6* KO E8.5 embryos. Most embryos used for this analysis were Type III mutants. Mean ± SEM. The statistical significance was calculated by Wilcoxon rank sum nonparametric test (n = 10, 8, 12, 12, 8, 9, 10, 9, 8, 13, 8, 10, 5, 7 in order of genes listed) (* at the *α* = 0.05 significance level, ** *α* = 0.01).

**Supplementary Figure 3.**
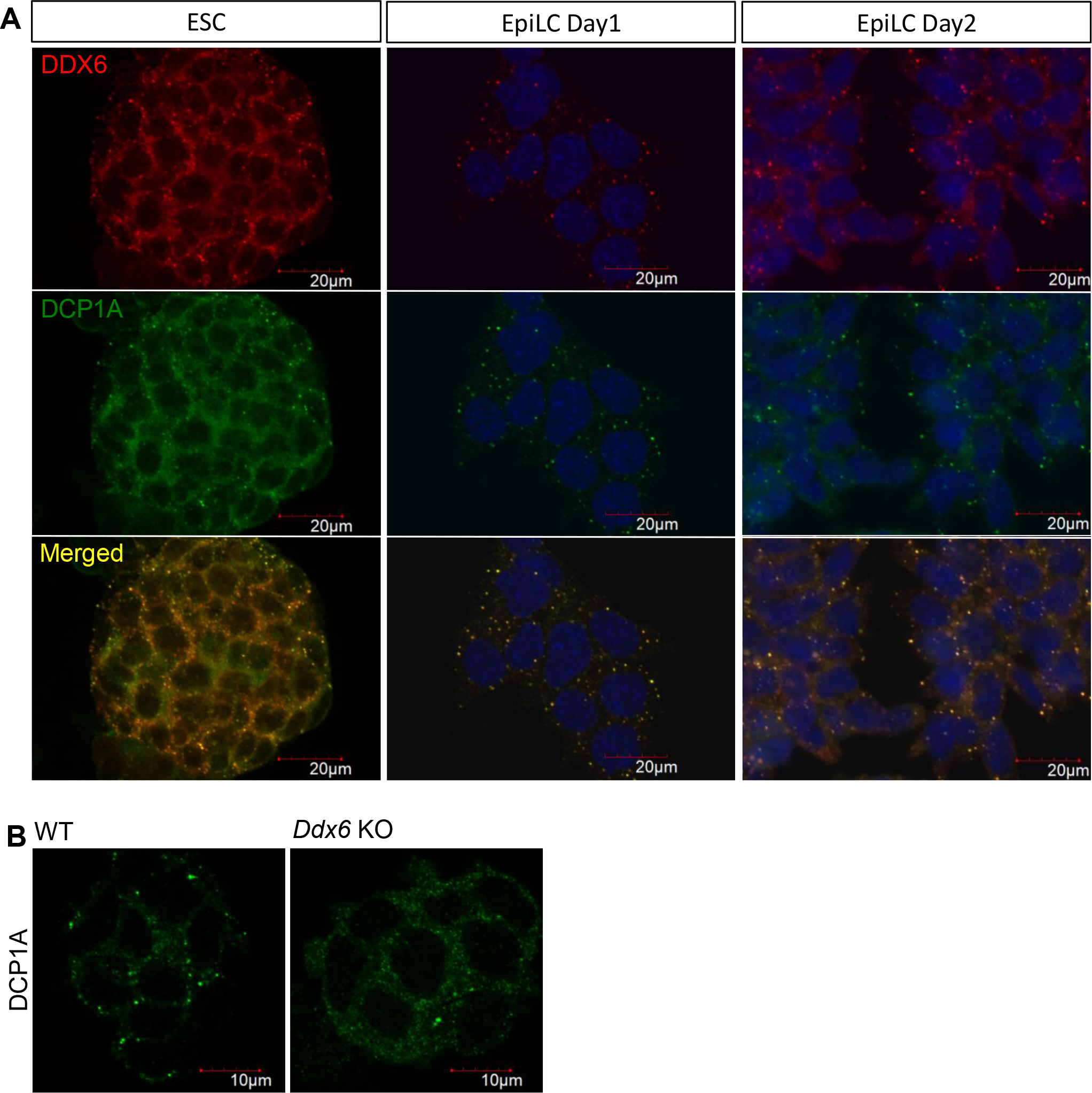
DDX6 and P-body expression in pluripotent cells. (A) ICC of DDX6 and a P-body marker DCP1A during ESC-to-EpiLC induction period. (B) Distinct granular P-bodies were disappeared in the absence of DDX6. ICC of DCP1A on ESCs.

**Supplementary Figure 4.**
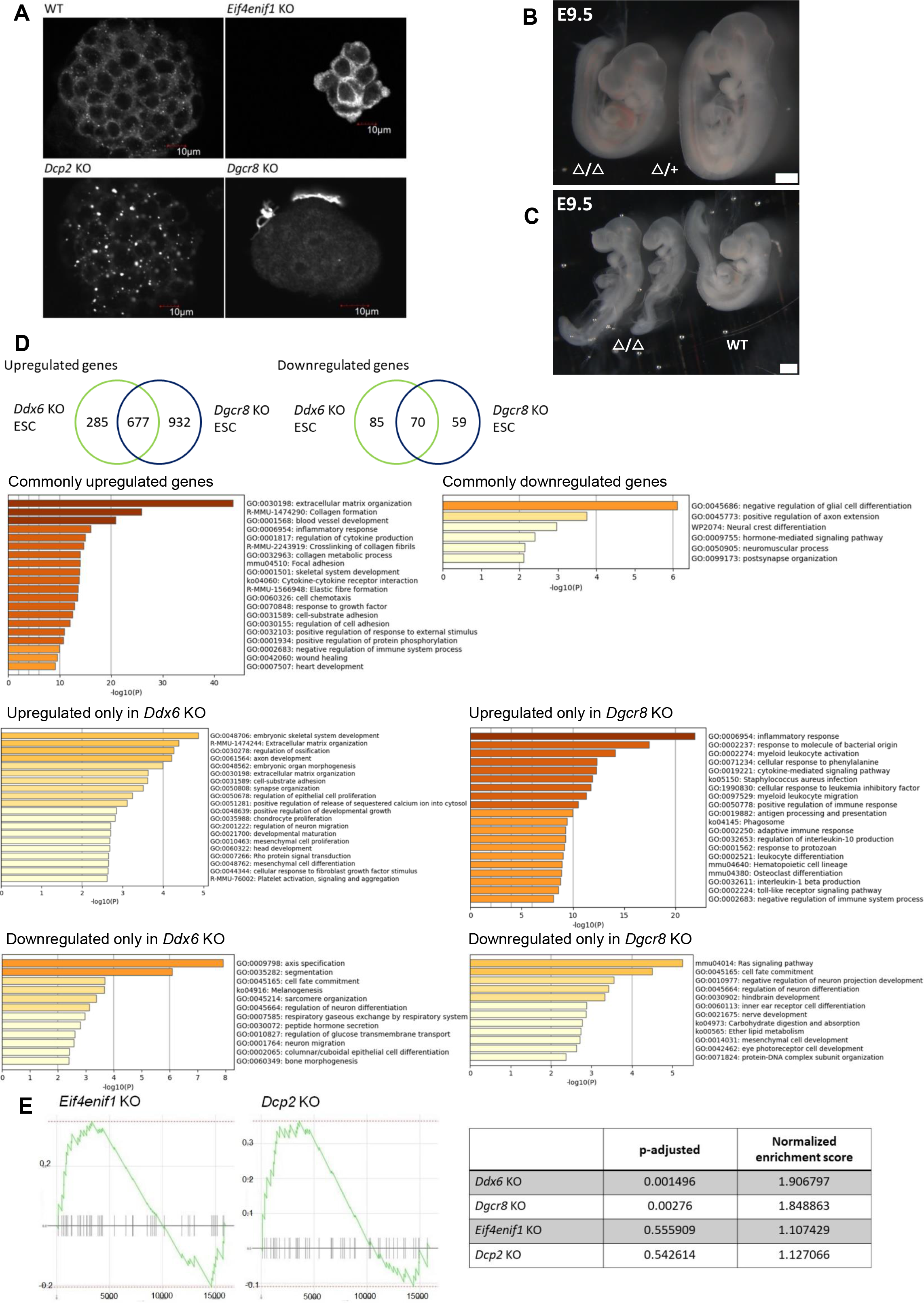
Examination of individual DDX6-mediated RNA regulator**y pathway.** (A) P-bodies in ESCs were affected by deletion of each gene. ICC of DDX6, a P-body marker, on ESCs. (B) E9.5 *Eif4enif1* KO embryo and a littermate control (scale: 500 μm, n = 7). (C) E9.5 *Dcp2* KO embryos and a littermate control (scale: 500 μm, n = 3). (D) Venn diagram showing the number of differentially expressed genes in *Ddx6* KO and *Dgcr8* KO ESCs. GO term enrichment analyses. (E) GSEA enrichment plot of the “negative regulation of BMP signaling pathway” gene set in *Eif4enif1* KO and *Dcp2* KO ESC as compared to WT ESC. Black bars represent the position of the genes that are belong to this gene set (n = 45) in the whole ranked gene list. The green line shows the overall distribution of this gene set (whether over-represented at the top (left) or bottom (right) of the ranked list of genes). The table includes the detailed results.

**Supplementary Figure 5.**
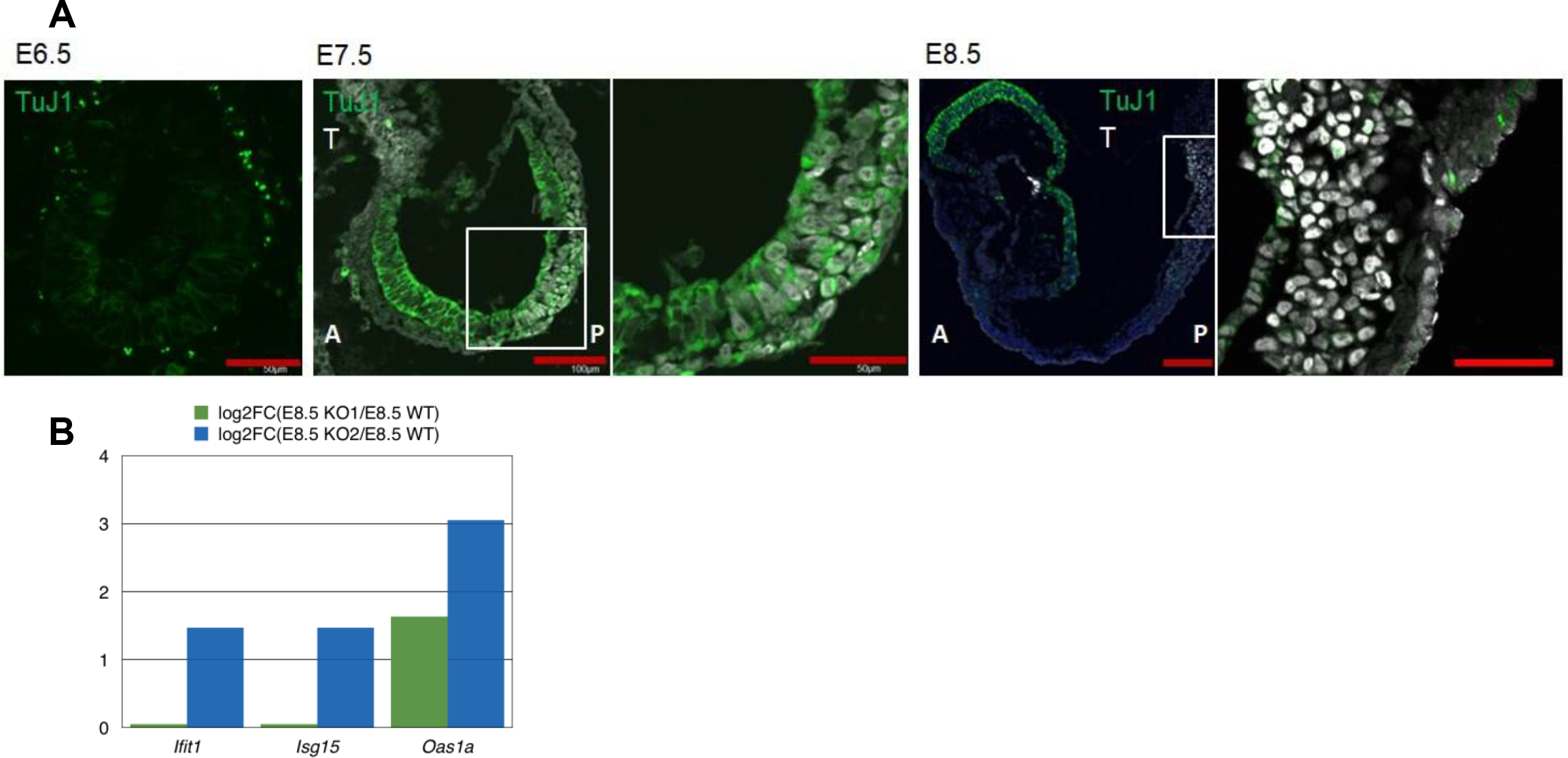
(A) TuJ1 expression in the epiblast. E6.5∼E8.5 embryo frozen section IHC for TuJ1 & T (50 μm for E6.5, n = 2; 100 μm for lower magnification, 50 μm for higher magnification, n = 3 for E7.5 & E8.5). (B) RNA-seq data comparing expression of interferon-stimulated genes (ISGs) in E8.5 *Ddx6* KO1 library (green) and *Ddx6* KO2 library (navy) with that in E8.5 WT.

**Table 1.** *Ddx6* knockout results in embryonic lethality by E11.5.

| Time point | +/+ | $\Delta$/+ | $\Delta$/$\Delta$ | Total |
| --- | --- | --- | --- | --- |
| E6.5 | 1 (7.1%) | 8 (57.1%) | 5 (35.7%) | 14 |
| E7.5 | 26 (41%) | 32 (51%) | 5 (8%) | 63 |
| E8.5 | 72 (33%) | 120 (55%) | 28 (13%) | 220 |
| E9.5 | 17 (23%) | 43 (59%) | 13 (18%) | 73 |
| E10.5 | 12 (42.9%) | 11 (39.3%) | 5 (17.9%) | 28 |
| E11.5 | 11 (44%) | 14 (56%) | 0 | 25 |
| E12.5 | 7 (30%) | 16 (70%) | 0 | 23 |

